# Evolution of Multicellularity Genes in the Lead Up to the Great Oxidation Event

**DOI:** 10.1101/2023.12.23.573081

**Authors:** Joanne S. Boden, Mercedes Nieves-Morión, Dennis J. Nürnberg, Sergio Arévalo, Enrique Flores, Patricia Sánchez-Baracaldo

## Abstract

Cyanobacteria are among the most morphologically diverse prokaryotic phyla on Earth. Their morphotypes range from unicellular to multicellular filaments, yet mechanisms underlying the evolution of filamentous morphologies remain unknown. Here, we implement phylogenomic, Bayesian molecular clock and gene-tree-species-tree reconciliation analyses to estimate when genes encoding cell-cell joining structures first evolved. We also characterise septal structures and measure intercellular communication rates in non-model and early-branching filamentous strains. Our results suggest that genes encoding septal proteins (namely *sepJ, sepI,* and *fraE*) and potentially pattern formation (*hetR*) evolved in the Neoarchaean ∼2.6-2.7 billion years (Ga) ago. Later, at the start of the Great Oxygenation Event ∼2.5 Ga, genes involved in cellular differentiation (namely *hetZ, patU3* and *hglK*) appeared. Our results predict that early-branching lineages like *Pseudanabaena* were capable of intercellular communication, but further innovations in cellular differentiation were needed to drive ecological expansion on a scale large enough to permanently oxygenate Earth’s atmosphere.

## Introduction

The Phylum Cyanobacteria emerged more than ∼3.3 Ga (*1-3*) when they diverged from their closest non-photosynthetic relatives, the Vampirovibrionia (*4, 5*). Nearly a billion years later, biological oxygen production by Cyanobacteria raised global atmospheric oxygen levels by several orders of magnitude during the Great Oxidation Event (GOE) of ∼2.45 to ∼ 2 Ga (*6, 7*). What caused this delay between the origin of photosynthetic Cyanobacteria and the GOE remains unclear (*8*). Insights have been provided by multiple lines of geochemical evidence which have helped characterise changes in the planet’s redox state (*7, 9, 10*), but less is known about the biological events that contributed to these geochemical changes (*11*).

A key driving factor may have been the ecological expansion of Cyanobacteria (*11*) in shallow water environments (*12*) prior to and during the GOE. This was likely driven by morphological innovations because phylogenetic and molecular clock analyses have found that early Cyanobacteria were simple unicellular organisms (*12-15*) and that filamentous cyanobacteria emerged toward the end of the Archean (*12-15*). The evolution of filaments was a key biological event (*16*) because it corresponded with an increase in lineage diversification (*13*), cell-to-cell communication (*13, 17*), and evolutionary innovations enabling the formation of thick laminated microbial mats and the colonization of shallow marine coastal habitats (*12*). However, a deeper biological understanding is required to unravel the interplay between life and environment on early Earth.

As a result of their evolutionary history, modern Cyanobacteria are some of the largest and most morphologically diverse phyla among prokaryotes (*18*). They encompass a range of unicellular and filamentous strains, some of which divide their labour between cells specialised for different functions. To distinguish between different morphologies, a classification system was developed which groups unicellular strains into sections I and II, undifferentiated filamentous strains into section III (although some produce hormogonia, motile filaments made of small cells) and differentiated filamentous strains into sections IV and V depending on their level of branching (*19*). Although it is tempting to assume that each section is evolutionarily unique, they are not. After filaments evolved in cyanobacteria shortly before the GOE, there were several backward transitions to unicellular morphology, resulting in a mixed phylogeny where unicellular strains are interspersed alongside the undifferentiated filaments of strains in section III in the tree of life (*13-15, 20*). Only the most complex of filaments, those with differentiated cells (sections IV and V) constitute a monophyletic group; the order Nostocales (see, e.g., (*1, 21-23*)).

The evolution of filaments was followed by the evolution of two of the most diverse and productive modern cyanobacterial clades: Macrocyanobacteria and Microcyanobacteria (*21*). These clades evolved after early-branching lineages, so can be considered as separate and distinct groups. Macrocyanobacteria is a monophyletic clade of lineages with large cell diameters of 3 μm to 50 μm (*21*), including taxa that are characteristic of marine microbial mats such as *Microcoleous* spp. (section III) as well as heterocyst-forming cyanobacteria of the Nostocales (sections IV and V) such as *Anabaena* spp. (*12, 24*). Heterocysts are micro-oxic cells specialised for N_2_ fixation that are found semi-regularly spaced in the filaments of these cyanobacteria under combined nitrogen deficiency (*16*). The Nostocales comprise a derived lineage within the Macrocyanobacteria that evolved cell differentiation and specialized labour, enabling them to occupy more diverse niches across terrestrial environments. In contrast, Microcyanobacteria is a monophyletic clade comprised of lineages with smaller cell diameters of <3 μm. Their appearance led to the evolution of some of the most abundant photosynthetic organisms on Earth (i.e., the marine *Synechococcus* and *Prochlorococcus*) (*25*).

Molecular biology studies, carried out mainly on *Anabaena* sp. PCC 7120 (hereafter *Anabaena*), have provided key insights into the structure and genomic mechanisms of achieving multicellularity. Their filaments consist of hundreds of cells which are maintained by a unique form of cell division characterized by the absence of peptidoglycan (PG) splitting and outer membrane invagination (*16*). Thus, within each filament, cells are enclosed by a continuous outer membrane and each cell is surrounded by PG (*16, 26*). Cell division is mediated by the divisome, a protein complex that includes proteins common to different bacteria and others that seem specific of cyanobacteria such as CyDiv (*26*).

Along the filament, intercellular communication takes place: Vegetative cells provide carbon compounds to heterocysts and heterocysts provide N_2_ fixation products to the vegetative cells (*27-29*). Large multi-protein complexes termed septal junctions are located at the intercellular septa where they connect neighbouring cells (*30-32*). These septal complexes are analogous to the “connexons” of gap junctions in large multicellular eukaryotes (*33, 34*). They traverse the septal PG layers through perforations termed nanopores and allow the transfer of metabolites such as glutamine, glutamate, alanine, β-aspartyl-arginine, and sucrose along the filament (*35-38*). Several septal proteins have been characterized in *Anabaena* that are part of the septal junctions, such as FraCD (*34, 39, 40*), or are needed for the formation of mature septa, such as HglK (*41*) or SepI (*42*). FraE, encoded in the *fraCDE* operon of *Anabaena* (*39*), is similar to ABC transporter permeases and is needed for heterocyst maturation (*40, 41*). Additionally, SepJ is a protein that stands out because of the strong phenotypic alterations of the *sepJ* mutants, including strong filament fragmentation, an early block in heterocyst differentaition, and a defect in nanopore formation and intercellular communication (*33, 34, 39, 40, 42-50*).

Vegetative cells in Nostocales differentiate into nitrogen-fixing heterocysts by regulating gene expression and establishing spatial patterning using gene products such as HetR, PatU3 and HetZ. Whilst PatU3 and HetZ control the frequency of heterocysts (*51, 52*), HetR is a transcriptional regulator needed for heterocyst differentiation to take place (*41, 53*). Surprisingly, some septal and heterocyst regulatory proteins are also present in filamentous non-heterocyst-forming cyanobacteria (section III), and in some unicellular strains (section II) such as *Gloeocapsopsis* sp. UTEX B3054 (*20, 42, 51, 54-57*).

Heterocyst-forming cyanobacteria like *Anabaena* (sections IV and V) have been evolving independently of undifferentiated filamentous cyanobacteria for hundreds of millions of years (*21*), but little research has addressed the evolution of genes involved in filament development in non-model organisms. Some studies suggest that genes encoding the septal protein SepJ evolved before early-branching strains including *Pseudanabaena* spp. diversified from heterocystous strains (*20*) and that the transcription factor HetR evolved in the most recent common ancestor of Microcyanobacteria (including *Synechococcus* sp. PCC 7335, a possible unicellular mutant of a filamentous cyanobacterium; (*40*)) and heterocystous strains (*54*). *Pseudanabaena* spp. were the earliest filamentous lineage to emerge (*13-15, 21*), so they offer the potential to provide insight into the timing of emergence of genes underpinning processes such as cell–cell adhesion and intercellular communication before cellular differentiation evolved in the Nostocales (*11*).

Here, we implement large-scale phylogenomic analyses, including Bayesian molecular clocks, to estimate when genetic mechanisms of creating filaments evolved and which cyanobacterial lineages diverged around the GOE. We also study the diversity of septal structures and kinetics of intercellular communication across major groups of cyanobacteria. To do this, we visualise the septal PG of non-heterocystous Macrocyanobacteria, Microcyanobacteria and *Pseudanabaena* by electron microscopy, and look for nanopores in their septal disks. We then implement FRAP (Fluorescence Recovery After Photobleaching) analyses using fluorescent tracer molecules (namely calcein and 5-carboxyfluorescein) to find out whether non-heterocystous cyanobacteria can engage in intercellular communication. The results show that early-diverging filamentous cyanobacteria contain less complex nanopores and are capable of intercellular molecular exchange, albeit at slower rates than those of the model and derived heterocystous strain *Anabaena*. Furthermore, comparative genomic analyses across 173 cyanobacterial genomes reveal that genes underlying intercellular communication and cell differentiation evolved shortly before and during the start of the Great Oxidation Event of 2.5 to 2.32 billion years ago.

## Results

### Evolution of Cyanobacteria

One hundred and seventy-three cyanobacterial genomes from strains encompassing a full variety of evolutionary histories, habitats, and morphologies were used to reconstruct the Cyanobacterial tree of life. The resulting phylogeny is broadly consistent with previous studies (*1, 2, 14, 21, 58-62*) and includes a variety of recently-sequenced genomes from (*60*). One of these, *Leptolyngbya* sp. FACHB-261 (also known as *Leptolyngbya* sp. FACHB-26 cyanobac1), is an early-branching filamentous strain related to *Pseudanabaena* ((*60, 63*) and UFBoot 100, Figure 1), so by incorporating it, the known diversity of early-branching filamentous lineages is expanded.

**Figure 1:**
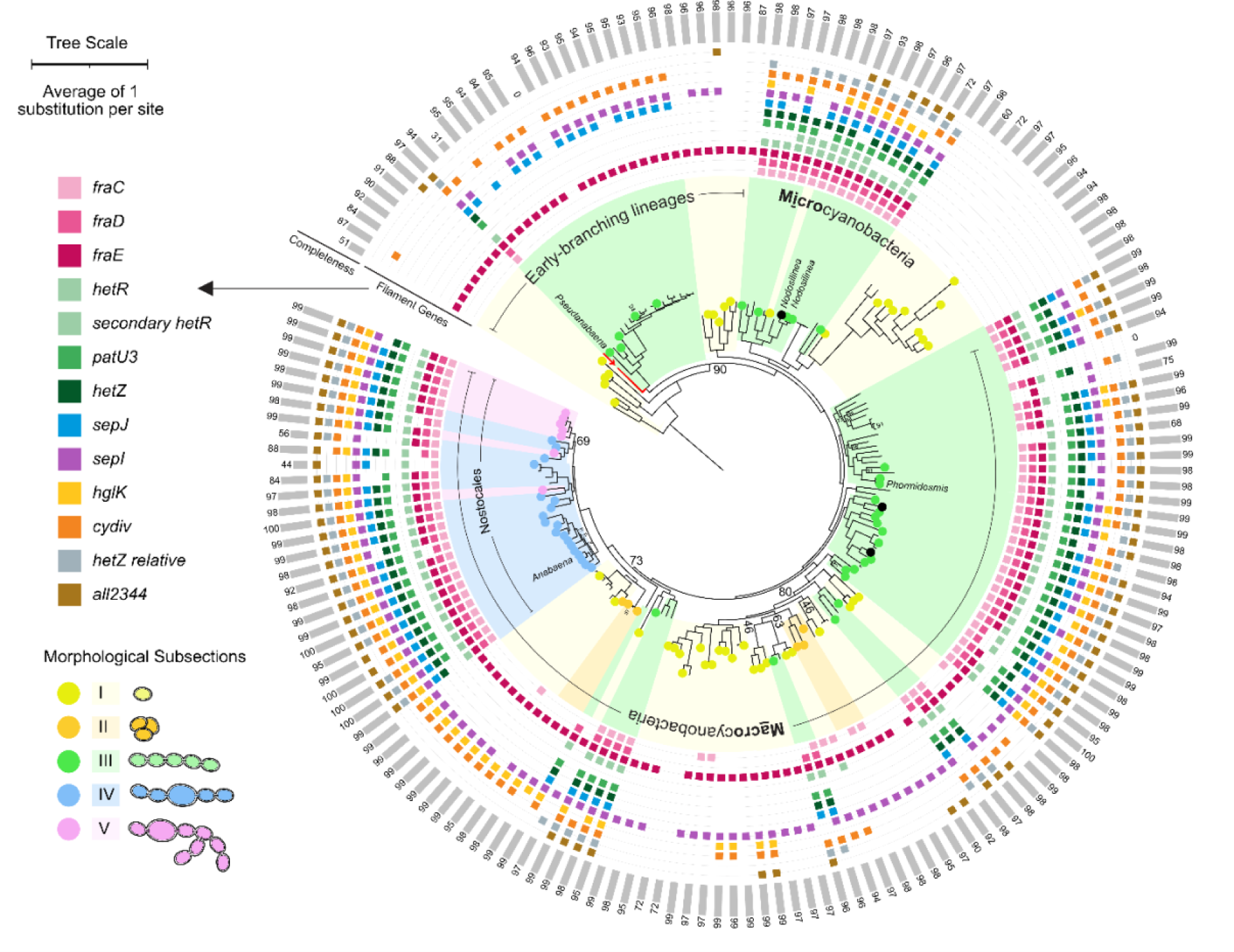
Distribution of genes underlying filamentous phenotypes in the Cyanobacterial tree of life. Homologs of genes encoding proteins required to form normal filaments (detailed in the main text) are marked with coloured squares. The evolutionary tree was estimated from SSU rRNA, LSU rRNA and 139 cyanobacterial proteins using maximum likelihood methodology implemented in IQ-TREE v1.6.1 and rooted with Vampirovibrionia. Numerical values on bifurcating branches represent ultrafast bootstrap support values < 95. Coloured circles on branch tips represent strains whose morphological section has previously been defined (Table S1). Black circles denote strains whose genomes were sequenced for this study. Grey bars and numbers on the outside of the tree represent percent completeness of the relevant strain’s genome, as measured by BUSCO version 3.0.2 using lineage data from Cyanobacteria (*66*). Two strains (namely *Pseudanabaena* sp. PCC 6903 and *Phormidium laminosum* Gom OH1pC11) whose genomes have not been sequenced were included based on ribosomal RNA because they showed potential for FRAP and septal disk visualisation, which later proved unsuccessful. Genus names represent strains whose septal structures were investigated. *Pseudanabaena* marks *Pseudanabaena* sp. PCC 7367, *Nodosilinea* marks *Nodosilinea* sp. PCC 9330 and *Nodosilinea* sp. LEGE 07298, *Phormidesmis* marks *Phormidesmis priestleyi* ULC007 and *Anabaena* marks *Anabaena* sp. PCC 7120. *Leptolyngbya* sp. FACHB-261 is highlighted in red with an arrow.

Unlike previous studies (e.g. (*21*)), which found that *Leptolyngbya* sp. PCC 7375 and *Leptolyngbya boryana* PCC 6406 belong to the LPP clade (*21, 59, 64*)) our phylogenetic reconstructions predict that LPP is a polyphyletic group (i.e. two well-supported clades with UFBoot 100, Figure 3B). This is consistent with recent phylogenomic analyses (*1, 62, 65*). We here name these clades Micro-LPP and Macro-LPP. MicroLPP are found within the Nodosilineales whilst MacroLPP are found within the Leptolyngbyales and Oculatellales (*65*). Two strains investigated for intercellular communication here (namely *Nodosilinea* sp. LEGE 07298 and *Nodosilinea* sp. PCC 9330 which was previously known as *Pseudanabaena)* are Micro-LPP.

**Figure 2:**
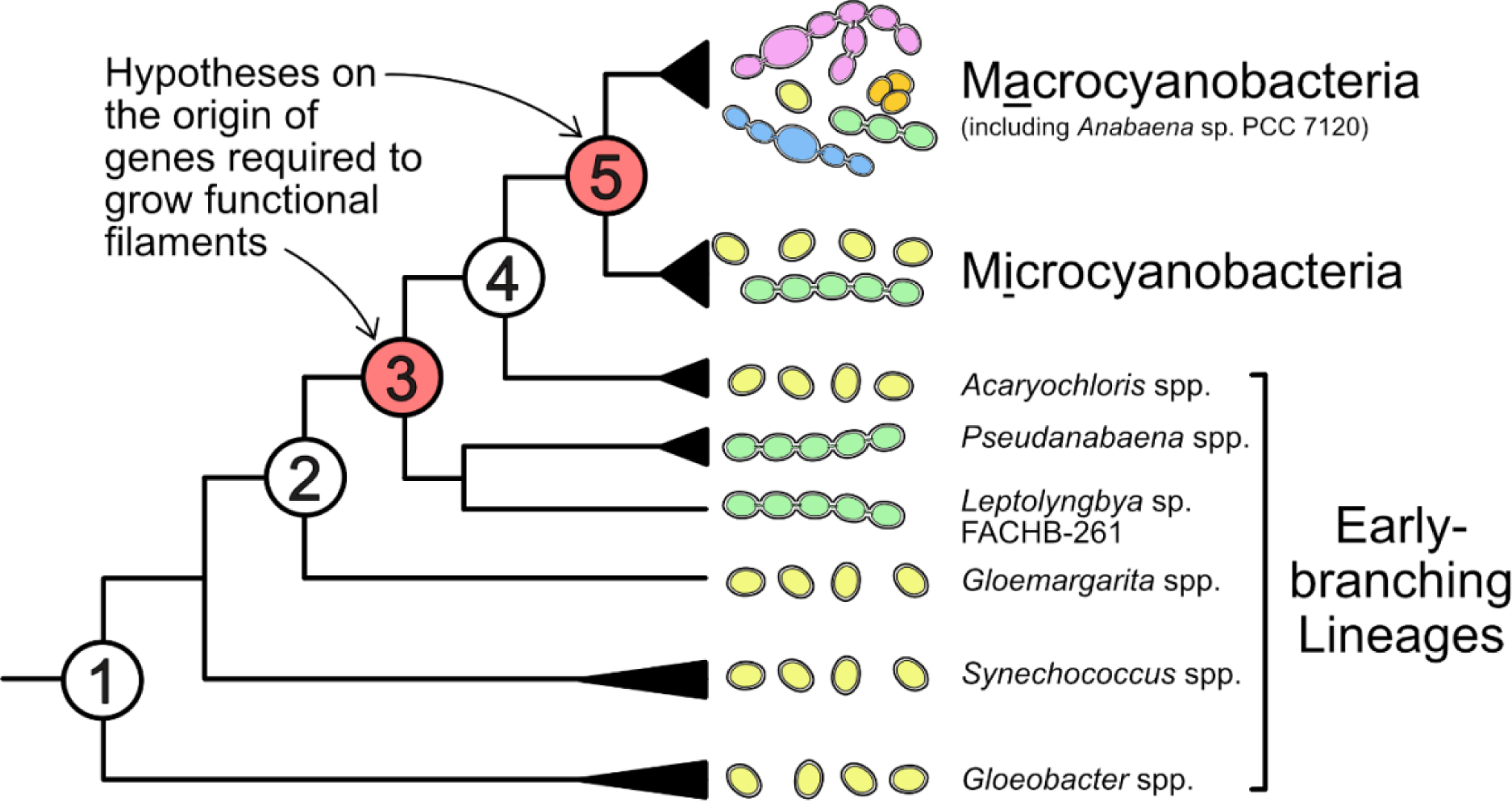
Two hypotheses on the origin of genetic mechanisms underlying filamentous morphology. Numbered circles mark nodes of interest.

**Figure 3:**
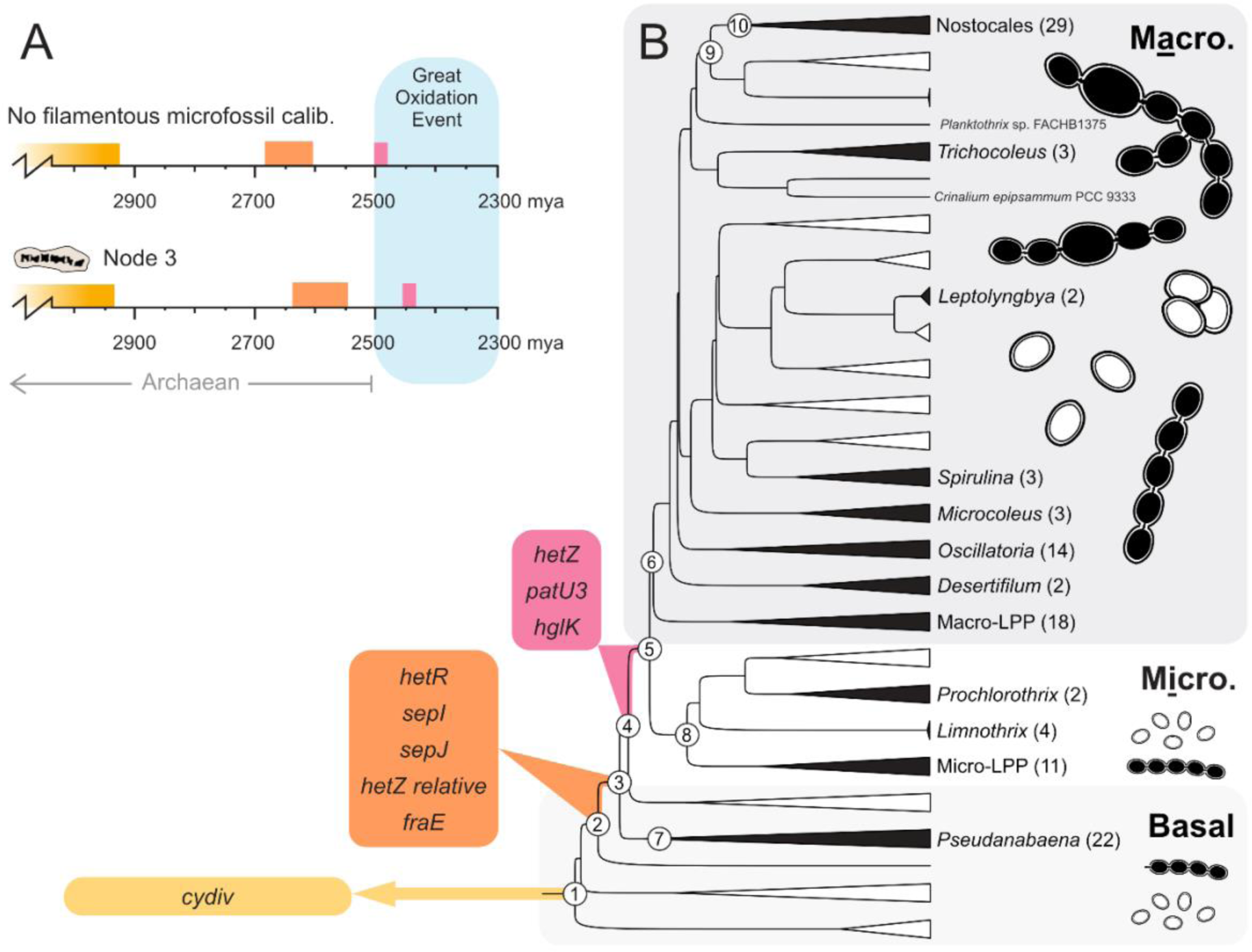
Estimated origin of genes associated with filament development in Cyanobacteria. Each box indicates which genes were present in the ancestral lineage of the same colour in the evolutionary tree. **A**. Molecular clock timeline estimates for genes highlighted in boxes of matching colour in B. The oldest microfossil of filamentous cyanobacteria is either not applied (top panel) or constrains the MRCA of all filamentous cyanobacteria (node 3) to more than 1.9 Ga (bottom panel). **B**. Black triangles and labelled lineages indicate predominantly filamentous clades, whereas white triangles and unlabelled lineages indicate predominantly unicellular clades. Some of the cartoon images are adapted from (*68*). The number of strains in each clade are presented in brackets alongside the clade names. Macro. refers to Macrocyanobacteria and Micro. to Microcyanobacteria.

### Comparative Genomics

All 173 cyanobacterial genomes were surveyed for genes encoding key septal proteins (i.e., *fraC, fraD, fraE, hglK, sepJ* and *sepI*) and regulators of cellular differentiation and pattern formation (i.e., *hetR*, *hetZ* and *patU3*). The function of these genes was originally determined in heterocystous strains such as *Anabaena* (see introduction for details), but we found homologues in other filamentous strains from morphological section III, whose cells cannot differentiate into heterocysts (Figure 1). These non-heterocystous filamentous strains are taxonomically diverse with cell sizes ranging from < 2 to > 5 μm in diameter (*21*). They include early-branching lineages (namely *Pseudanabaena* spp. and *Leptolyngbya* sp. FACHB-261), Microcyanobacteria (including *Prochlorothrix hollandica, Limnothrix spp.* and *Nodosilinea* spp.,) and Macrocyanobacteria (including *Phormidesmis* spp., *Trichodesmium* spp., *Oscillatoria* spp., *Spirulina* spp., *Crinalium epipsammum* and *Microcoleus* spp.). Among these strains of section III, early-branching lineages of *Pseudanabaena* spp. are unique because they lack genes encoding septal junction proteins (namely *fraC* and *fraD* and *hglK*) involved in intercellular communication and the formation of nanopores in *Anabaena* (*33, 39, 40, 50*). They also lack genes involved in heterocyst differentiation (namely *hetR*) and pattern formation (namely *hetZ*, *patU3*; (*52*) (Figure 1). These genes are present, however, in *Leptolyngbya* sp. FACHB-261 (Figure 1).

In contrast, most unicellular cyanobacteria lack *sepJ*, *fraC, fraD, hetR, hetZ* and *patU3,* but many maintain a permease (encoded by *fraE*), a cyanobacterial division protein originally described in filamentous cyanobacteria (encoded by *cyDiv*), a pentapeptide-repeat-containing protein (encoded by *hglK*) and a septal protein (encoded by *sepI*) which are required to maintain and develop filaments in *Anabaena* (Figure 1). On the other hand, filamentous cyanobacteria with branches (and therefore belonging to section V, e.g., *Fischerella* spp., *Mastigocoleus testarum* and *Mastigocladopsis repens* PCC 10914) are not monophyletic (Figure 1) because some have evolved from ancestors which also gave rise to non-branching strains (from section IV, Figure 1).

### Timing the Origin of Genes Underlying Filamentous Morphology

Relaxed molecular clock analyses were implemented to estimate when genetic mechanisms of creating filaments emerged. Timing of origin for these genes was estimated by testing two hypotheses (Figure 2) using information derived from reconciliations between the evolutionary history of each gene (Figures S1-10) and the Cyanobacterial tree of life (i.e., genome phylogeny; Figure 1). These hypotheses state that either the gene arose in the most recent common ancestor (MRCA) of all filamentous cyanobacteria (Node 3), or that the gene arose in the MRCA of filamentous Macro- and Micro-cyanobacteria, but not early-branching lineages including *Pseudanabaena* spp. and *Leptolyngbya* sp. FACHB-261 (Node 5).

Phylogenetic evidence suggests that key genes for maturation of septal structures in filaments appeared in ancestral cyanobacteria in the lead-up-to the GOE. These include *sepJ* and *sepI* which encode septal proteins that are needed to localise septal junctions during cell division. Also, the permease encoded by *fraE* is included, because the homologs present in early-branching strains are more basal than those of later-branching Micro- and Macro-cyanobacteria (Figures S3, S4 and S6) and therefore consistent with vertical inheritance from the MRCA of all filamentous species (Figure 3, node 3). As a result, *sepI, sepJ* and *fraE* must have first emerged somewhere on the lineage between nodes two and three, which existed in the Neoarchaean from ∼2.70 to ∼2.60 Ga (95% credibility intervals span 2.875 to 2.384 Ga). An additional gene, *hetR* encoding the heterocyst master regulator, may also have appeared on this lineage despite its absence from *Pseudanabaena* spp. This is because a different early-branching strain (namely *Leptolyngbya* sp. FACHB-261) encodes a *hetR* homolog which is basal to those of other *hetR* (Figure S2, Posterior Probability 85), consistent with vertical inheritance from node three and a subsequent loss of the gene in *Pseudanabaena* spp. (Figure S12).

Further genes guiding the development of differentiated cells (namely *hetZ* and *patU3*) and architecture of the intercellular septa (*hglK*) show evolutionary patterns which are consistent with vertical inheritance from the MRCA of Macro- and Micro-cyanobacteria (Figure 3, node 5, Figures S5, S7 and S8). Therefore, they must have evolved somewhere on the lineage between nodes four and five, which existed during the GOE from ∼2.52 to ∼2.47 Ga (95% credibility intervals span 2.700 to 2.289 Ga) (Figure 3, Table 1). Results for *fraC* and *fraD* were inconclusive because their phylogenies include many polytomies and low support values which hinder inferences about their origins (Figures S9-10).

**Table 1:**
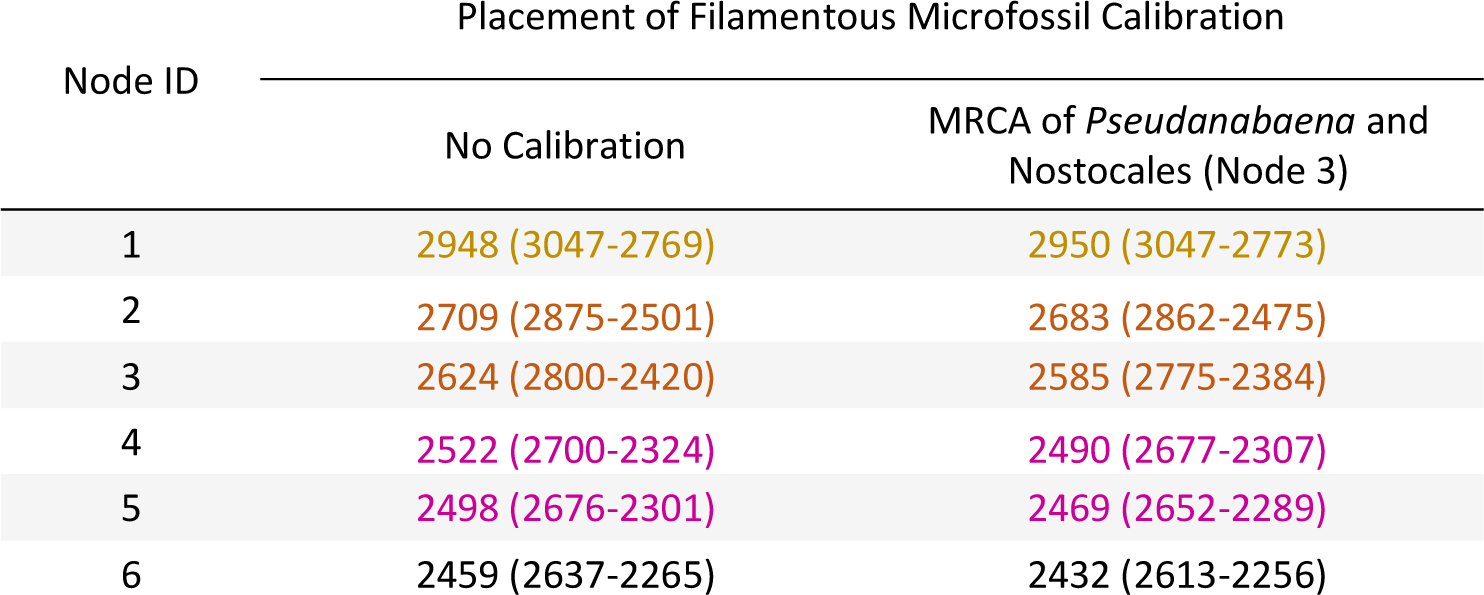
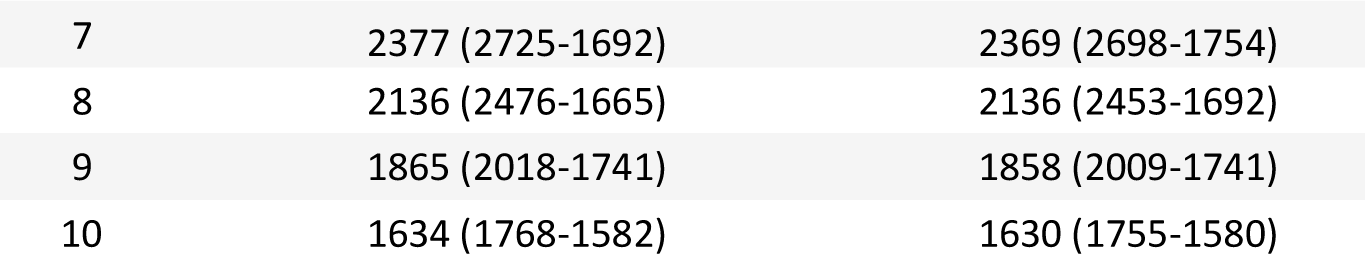
Posterior age estimates using a Bayesian approach. Mean divergence times with 95% credibility intervals in brackets. Node IDs refer to key diversification events presented within numbered circles in Figure 3B with coloured text matching that of Figure 3.

| Node ID | Placement of Filamentous Microfossil Calibration |  |
| --- | --- | --- |
|  | No Calibration | MRCA of *Pseudanabaena* and Nostocales (Node 3) |
| 1 | 2948 (3047-2769) | 2950 (3047-2773) |
| 2 | 2709 (2875-2501) | 2683 (2862-2475) |
| 3 | 2624 (2800-2420) | 2585 (2775-2384) |
| 4 | 2522 (2700-2324) | 2490 (2677-2307) |
| 5 | 2498 (2676-2301) | 2469 (2652-2289) |
| 6 | 2459 (2637-2265) | 2432 (2613-2256) |
| 7 | 2377 (2725-1692) | 2369 (2698-1754) |
| 8 | 2136 (2476-1665) | 2136 (2453-1692) |
| 9 | 1865 (2018-1741) | 1858 (2009-1741) |
| 10 | 1634 (1768-1582) | 1630 (1755-1580) |

The gene *cyDiv*, which is involved in cell division to maintain straight filaments (*26*) probably pre-dates these genes involved in cell differentiation and the production of septal junctions because it is encoded in the genomes of many unicellular strains, including one of the earliest-branching cyanobacteria, *Gloeobacter violaceus* PCC 7421. As a result, it exhibits an evolutionary trajectory indicative of vertical inheritance from the most recent common ancestor of all Cyanobacteria (Figure 3, node 1, Figure S1). This observation places a younger bound on the origin of *cyDiv* in the Mesoarchaean, 2.95 Ga (95% credibility intervals span 3.047 to 2.769 Ga) (Figure 3, Table 1).

Many unicellular strains (section II and some of section I) evolved from a filamentous ancestor at node five which is predicted to have harboured multiple genes encoding components of septal junctions and cell-cell patterning, so they must have lost multiple genes as well as their filamentous morphology. These include *Synechocystis* sp. PCC 6803 and *Synechococcus* sp. PCC 7002 (model strains in many experimental analyses) as well as *Dactylococcopsis salina* PCC 8305, *Geminocystis hermanii* PCC 6308*, Pleurocapsa* sp. PCC 7319 and *Prochloron didemni* (symbionts, (*67*)).

### Calibrating the Origin of Filamentous Cyanobacteria

The microfossil-based calibration which constrains the origin of filamentous species at node 3 (the MRCA of all filamentous species, including early-branching lineages) to more than 1.9 Ga has very little impact on the estimated emergence of multicellularity genes in our molecular clock. This is in comparison to analyses which do not apply the calibration on any node in the tree (Table 1).

### Genetic Context

To assess whether gene clusters are conserved across different lineages, we studied the surrounding genomic loci of septal structure-related genes. Genes encoding the septal junction proteins FraC and FraD are found on the same locus as a gene encoding an ABC-type permease (namely *fraE*) in all filamentous cyanobacteria except the early-branching *Pseudanabaena*, which lack *fraCD* (Table 2). Similarly, we found that less than 3,000 nucleotides separate *hetR* from *sepJ* in the genomes of multiple heterocyst-forming filamentous strains from morphological sections IV and V. The *hetR* and *sepJ* genes of filamentous strains with undifferentiated cells in morphological section III are not co-localised in gene clusters like this. Therefore, the clustering of genes guiding the formation of heterocysts and septal-related structures is a unique feature of filamentous strains with differentiated cells (sections IV and V; Nostocales). In contrast, the clustering of genes encoding septal junction proteins is widespread in all filamentous strains containing such genes.

**Table 2:** Genomic localisation of genes associated with filament development in Cyanobacteria from morphological sections III, IV and V. Each row indicates whether a group of genes (either *fraC*, *fraD* and *fraE* or *hetZ* and *patU3* or *hetR* and *sepJ*) are co-localised in the genomes of a phylogenetically distinct group of filamentous cyanobacteria. Coloured cells indicate that the relevant gene is present and adjacent to another described in the table under the same locus heading. In these cases, the percentage describes the proportion of strains where the gene is in that position. White cells represent that the gene is either absent or not adjacent to another described under the same locus heading. If the gene is present in a different locus, a black ‘P’ is presented, whereas if the gene is absent, only a percentage is shown, indicating the proportion of genomes without the gene. Proportions alongside a black ‘P’ indicate the proportion of genomes where the genes are present in a different location. If some strains have co-localised genes, but others do not, the pattern of most strains in the relevant clade is reported (detailed breakdown presented in Table S2). The position from left to right is not intended to define directionality. For example, *fraE* is near *fraD* in the genome of *Leptolyngbya* sp. FACHB261 but could be upstream or downstream. The column entitled ‘proportion’ describes the number of genomes with the pattern presented. * Each gene is separated by < 1,000 nucleotides. ** each gene is separated by < 3,000 nucleotides. P, present in a different part of the genome. M. S. Morphological Section

| Species Tree | M. S. | Clade Name | Locus One * |  |  | Locus Two * |  | Locus Three ** |  | Sample Size (n) | Proportion with presented pattern |
| --- | --- | --- | --- | --- | --- | --- | --- | --- | --- | --- | --- |
|  |  |  | *fraC* | *fraD* | *fraE* | *hetZ* | *patU3* | *hetR* | *sepJ* |  |  |
|  | III | *Leptolyngbya* sp. FACHB261 | 100% | 100% | 100% | 100% | 100% | P (100%) | P (100%) | 1 | 100% |
|  | III | *Pseudanabaena* | 100% | 100% | P (90%) | 100% | 100% | 100% | P (80%) | 20 | 75% |
|  | III | Micro-LPP | 100% | 100% | 75% | 88% | 88% | P (75%) | P (75%) | 8 | 63% |
|  | III | *Prochlorothrix* | 100% | 100% | 50% | 100% | 100% | P (100%) | P (50%) | 2 | 50% |
|  | III | *Limnathrix* | 100% | 100% | 100% | 100% | 100% | P (100%) | P (100%) | 4 | 100% |
|  | III | Macro-LPP | 71% | 88% | 59% | 94% | 94% | P (82%) | P (65%) | 17 | 35% |
|  | III | *Desertifilum* | 100% | 50% | 100% | 100% | 100% | P (100%) | P (100%) | 2 | 50% |
|  | III | *Oscillatoria* | 79% | 86% | 71% | 93% | 86% | P (100%) | P (93%) | 14 | 57% |
|  | III | *Microcoleus* | 100% | 100% | 67% | 100% | 100% | P (67%) | P (67%) | 3 | 67% |
|  | III | *Spirulina* | 100% | 100% | P (67%) | 100% | 100% | P (100%) | P (67%) | 3 | 33% |
|  | III | *Limnathrix rosea* and *Leptolyngbya* sp. PCC 7376 | P (50%) | P (100%) | P (100%) | 100% | 100% | P (100%) | P (100%) | 2 | 50% |
|  | III | *Crinalium epipsammum* PCC 9333 | 100% | 100% | 100% | 100% | 100% | P (100%) | P (100%) | 1 | 100% |
|  | III | *Trichocoleus* | 100% | 100% | 67% | 100% | 100% | P (100%) | P (100%) | 3 | 67% |
|  | III | *Phormidium* | 50% | 100% | 100% | 50% | 100% | P (100%) | P (100%) | 2 | 50% |
|  | IV and V | Nostocales | 100% | 97% | 79% | 90% | 86% | 100% | 62% | 29 | 55% |

### Novel Genes Warrant Further Study

Whilst investigating the origin and evolution of genes underlying filamentous morphology, BLAST analyses revealed three genes of unknown function that may provide further insight into the maintenance and evolution of filaments. These are: (i) a paralog of *hetZ* named the ‘*hetZ relative*’ on NCBI, which is unique to filamentous strains and emerged at the beginning of the GOE with *sepJ, sepI* and *hetR*, but currently has no known function; (ii) a paralog of *hetR* termed ‘*secondary hetR*’ is unique to filamentous Microcyanobacteria, but has no known function; (iii) a hypothetical gene (named *all2344* in *Anabaena*) was found to be associated with filamentous cyanobacteria in a previous study (*57*), but here we find that it emerged in the same lineage as *hetZ*, *patU3* and *hglK* and is characteristic of filamentous Macro- and Micro-cyanobacteria, but not *Pseudanabaena* spp..

### Intercellular communication and septal structures

Our phylogenetic analyses show that genes guiding the development of filaments and cell differentiation in Nostocales are also present in diverse Macro- and Micro-cyanobacteria as well as early-branching *Pseudanabaena* spp.. For example, *sepJ* is widespread in all filamentous cyanobacteria, whereas *fraC* and *fraD* are present in Macro- and Micro-cyanobacteria but not in *Pseudanabaena* spp.. To test directly for intercellular communication in filamentous cyanobacteria other than the Nostocales, we studied in selected strains intercellular transfer of two fluorescent markers, calcein and 5-carboxyfluorescein (5- CF) (622.5 Da and 376 Da, respectively, both negatively charged but to different extents (*69*)), by means of FRAP analyses (*40, 44*). In these analyses, hydrophobic, non-fluorescent forms of the markers diffuse into the cytoplasm of the cells where they are processed to hydrophilic and fluorescent forms. The fluorescence is bleached out in a single cell by an increased laser intensity, and recovery of fluorescence resulting from transfer of the marker from adjacent cells by diffusion is followed for a short time. The recovery rate constant (*R*) describes how quickly each marker is transferred from adjacent cells through their septal pores (*40*).

FRAP analysis of an early-branching filamentous strain, *Pseudanabaena* sp. PCC 7367, and the filamentous Microcyanobacterium *Nodosilinea* sp. PCC 9330, confirm fluorescence recovery of 5-CF in bleached cells (Figure 4), despite the lack of the septal junction genes *fraC* and *fraD* in *Pseudanabaena* sp. PCC 7367 (Figure 1). Intercellular transfer of at least one marker was also observed in another filamentous Microcyanobacterium, *Nodosilinea* sp. LEGE 07298, and a Macrocyanobacterium which lacks differentiated cells, *Phormidesmis priestleyi* ULC007, albeit at slower rates than those of the model strain, *Anabaena* (Table 3). Notably, with some strains and markers, two populations of cells could be discerned regarding intercellular communication: Communicating cells (typically, *R* values > 0.01 s^-1^) and non-communicating cells. The latter exhibit very low or null *R* values, suggesting regulation of the process as initially described in *Anabaena* (*34, 70*). These two populations were found in the Macrocyanobacterium *Phormidesmis priestleyi* ULC 007 with calcein; in the two Microcyanobacteria, *Nodosilinea* sp. PCC 9330 with 5-CF and *Nodosilinea* sp. LEGE 07298 with calcein and 5-CF; and in the early-branching cyanobacterium *Pseudanabaena* sp. PCC 7367 with 5-CF. Therefore, our results indicate that regulated intercellular molecular exchange occurs beyond the Nostocales, including Macro-, Micro- and early-branching cyanobacteria. Of note, *Pseudanabaena* sp. PCC 7367 was able to transfer 5-CF but not calcein (Figure 4).

**Figure 4.**
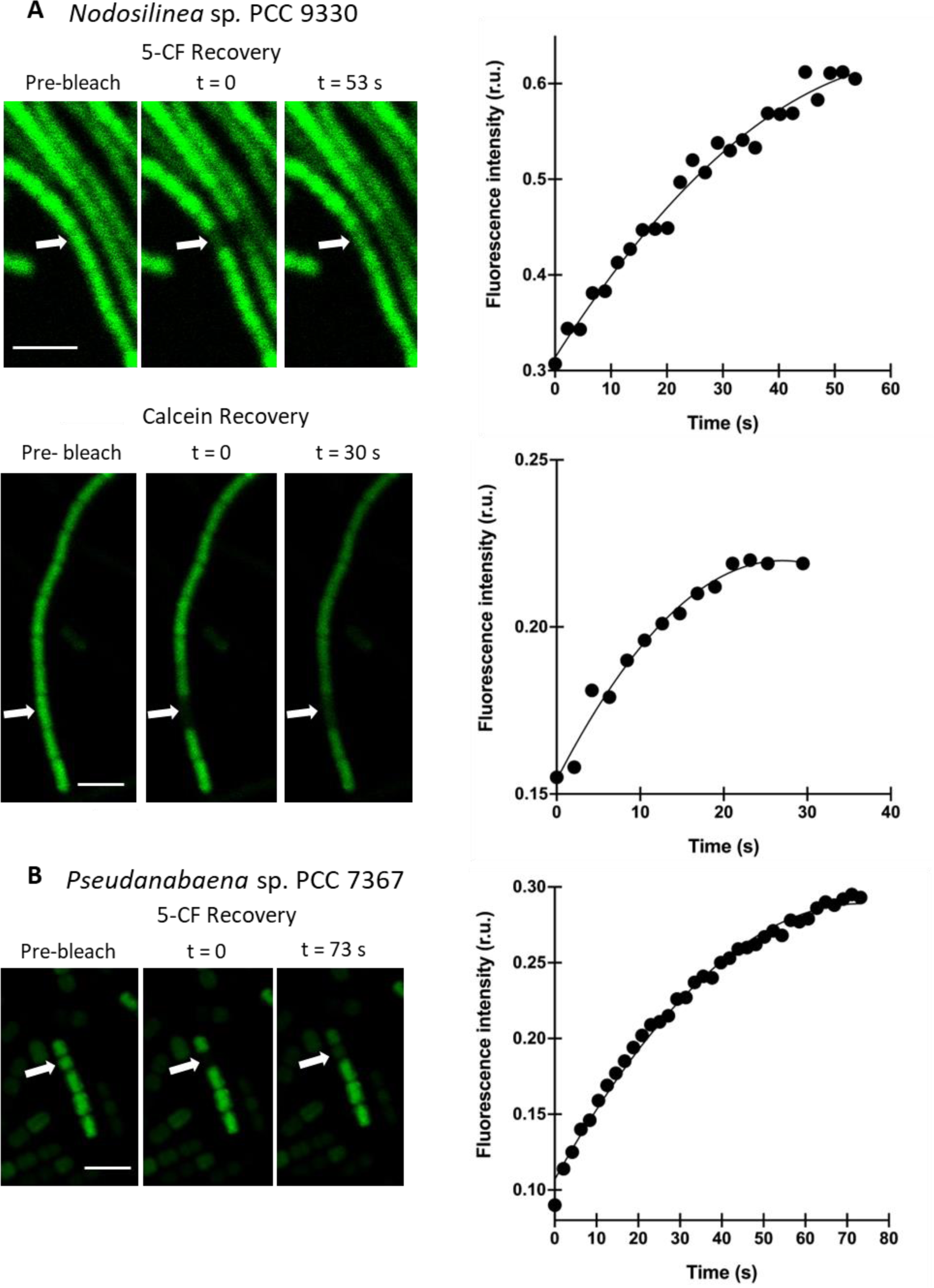
Intercellular molecular transfer in *Nodosilinea* sp. PCC 9330 (A) and *Pseudanabaena* sp. PCC 7367 (B). Examples of FRAP analysis for filaments from independent cultures of *Nodosilinea* sp. PCC 9330 (A) and one filament of *Pseudanabaena* sp. PCC 7367 (B) grown in BG11 and ASW medium, respectively. Filaments were labelled with 5-carboxyfluorescein (5-CF) or calcein as indicated, and subjected to FRAP analysis. No calcein recovery was observed for *Pseudanabaena* sp. PCC 7367. Images are shown at the times indicated. The plots show the recovery of fluorescence intensity in the bleached cell over time after the bleaching event (note the different scales). The line is an exponential curve fitted to the data points as described in the Materials and Methods. Scale bars, 5 µm. Note the different magnifications in the 5-CF and calcein experiments for PCC 9330.

**Table 3.** FRAP analysis and septal structures in different filamentous cyanobacteria. FRAP analysis: Rate constants (*R*) of calcein and 5-carboxyfluorescein (5-CF) recovery in the indicated strains and, for comparison, the model cyanobacterium *Anabaena* are shown. Note that two populations of cells, communicating cells (*R*_high_, > 0.01 s^-1^) and non-communicating cells (*R*_low_, < 0.01 s^-1^), were observed for transfer of a marker in some strains. The presence of communicating and noncommunicating cells in cyanobacterial filaments has previously been described and discussed for *Anabaena* (*41*). Data for communicating cells of *Anabaena* are from (*69*). Septal structures: Data from septal structures of the indicated strains and the model cyanobacterium *Anabaena* for comparison. Mean and standard deviation of the number and diameter of nanopores or central pores for each strain analysed are shown. Data for *Anabaena* are for vegetative cells, taken from (*70*) and Arévalo and Flores (2021). Calcein FRAP could also be determined for *Prochlorothrix hollandica* PCC 9006 (Microcyanobacteria), *R =* 0.118 ± 0.033 s^-1^ (n=19)..

| Strain | Phylogenetic group | FRAP recovery constant ($R$)<br>Mean ± SD nm (n) | | Septal disks and nanopores<br>Mean ± SD nm (n) | | |
| --- | --- | --- | --- | --- | --- | --- |
| | | Calcein<br>($\text{s}^{-1}$) | 5-Carboxyfluorescein<br>($\text{s}^{-1}$) | Septal disk<br>diameter (nm) | Number of<br>nanopores or<br>central pore*/disk | Nanopore or<br>central pore<br>diameter (nm) |
| *Anabaena* sp.<br>PCC 7120 | Macro | $R_{\text{high}}$, $0.097 \pm 0.023$ (18) | $R_{\text{high}}$, $0.080 \pm 0.013$ (37) | $850 \pm 53$ (7) | $41.67 \pm 21.85$ (21) | $17.06 \pm 4.13$ (131) |
| *Phormidesmis*<br>*priestleyi*<br>ULC 007 | Macro | $R_{\text{high}}$, $0.020 \pm 0.010$ (21)<br>$R_{\text{low}}$, $0.003 \pm 0.003$ (5) | No staining | $2003 \pm 230$ (27) | $7.25 \pm 4.60$ (27) | $13.83 \pm 3.37$ (46) |
| *Nodosilinea* sp.<br>PCC 9330 | Micro | $R_{\text{high}}$, $0.062 \pm 0.032$ (10)<br>$R_{\text{low}}$, not observed | $R_{\text{high}}$, $0.016 \pm 0.006$ (9)<br>$R_{\text{low}}$, $0.005 \pm 0.005$ (10) | $466 \pm 74$ (18) | $2.80 \pm 1.50$ (18) | $22.30 \pm 4.80$ (49) |
| *Nodosilinea* sp.<br>LEGE 07298 | Micro | $R_{\text{high}}$, $0.052 \pm 0.039$ (13)<br>$R_{\text{low}}$, $0.001 \pm 0.001$ (2) | $R_{\text{high}}$, $0.032 \pm 0.012$ (9)<br>$R_{\text{low}}$, $0.000$ (20) | $601 \pm 66$ (11) | $5.55 \pm 2.80$ (11) | $20.36 \pm 3.17$ (48) |
| *Pseudanabaena*<br>sp. PCC 7367 | Basal | 0.000 (15) | $R_{\text{high}}$, $0.014 \pm 0.003$ (11)<br>$R_{\text{low}}$, $0.002 \pm 0.003$ (18) | $324 \pm 136$ (40) | *1 (35); 0 (5) | $40.10 \pm 20.0$ (35) |

In Nostocales, intercellular molecular transfer takes place through septal junctions, which are proteinaceous structures that traverse the septal peptidoglycan through holes termed nanopores (*34*) (*70*). Nanopores can be conveniently visualized in septal peptidoglycan disks that are observed in murein sacculi preparations of filamentous cyanobacteria (*71*). We therefore looked for the presence of nanopores in septal disks from some of the strains for which we observed intercellular molecular exchange. Nanopores were observed in *Phormidesmis priestleyi* ULC 007, the two *Nodosilinea* strains, PCC 9330 and LEGE 07298, and *Pseudanabaena* sp. PCC 7376 (Figure 5). Interestingly, the septal disks of *Phormidesmis priestleyi* ULC 007 (Macrocyanobacteria) were large, about 2.36-fold larger than those of *Anabaena*, whereas the disks of *Nodosilinea* spp. (Microcyanobacteria) were smaller than those of *Anabaena*, and the septal disks of *Pseudanabaena* sp. PCC 7376 (an early-branching cyanobacterium) were even smaller (Table 3). The number of nanopores in *Phormidesmis priestleyi* ULC 007 and the two *Nodosilinea* strains was about 17% and 7-13%, respectively, of the number found in *Anabaena*, although nanopore size was not very different (about ± 20% the size in *Anabaena*). In contrast, *Pseudanabaena* sp. PCC 7376 had a single hole at the centre of the disk that was larger than the nanopores of any other cyanobacterium; we denote it a “central pore”. Together, FRAP analysis and visualization of septal disk nanopores indicate that structures for intercellular molecular exchange are present in filamentous Macrocyanobacteria, Microcyanobacteria and early-branching *Pseudanabaena*, although the structures of *Pseudanabaena* sp. PCC 7376 may have unique characteristics.

**Figure 5.**
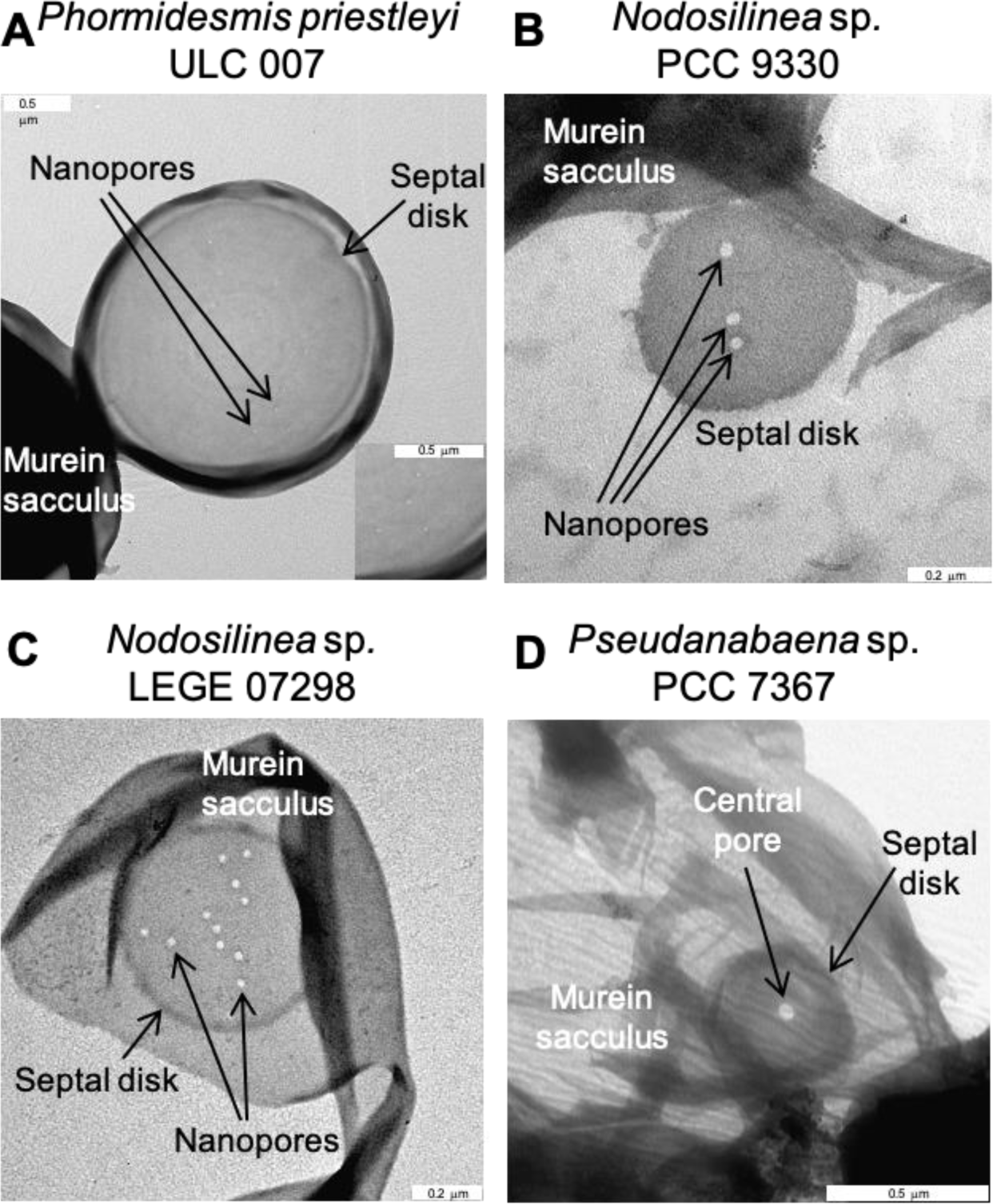
Septal structures in *Phormidesmis priestleyi* ULC 007 (A), *Nodosilinea* sp. PCC 9330 (B), *Nodosilinea* sp. LEGE 07298 (C) and *Pseudanabaena* sp. PCC 7367 (D). The peptidoglycan was isolated and visualized by transmission electron microscopy as described in Materials and Methods. (A, B, C) Murein sacculus, septal disks and nanopores are shown. (D) Murein sacculus, septal disk and a central pore are shown.

## Discussion

### Timing of Evolutionary Events

The transition from unicellular to filamentous morphology changed the ecological landscape during the Neoarchean and Paleoproterozoic by facilitating the formation of microbial mats and ecological expansion of oxygenic primary producers (*13*). Here, we provide insight into how filamentous cyanobacteria emerged by dating the origin of genes required for intercellular communication and cellular differentiation. Our analyses are consistent with trait evolution studies estimating that the earliest cyanobacteria (i.e., LCA of Cyanobacteria) were unicellular (*12, 14, 21, 72*) because the earliest-branching taxa lacked many of the genes needed for cell-to-cell communication (Figure 3).

The first septal proteins evolved during the late Archean between 2.7 and 2.6 Ga (Figure 3, Table 1), which corresponds with trait evolution studies estimating the origin of filaments in the lead up to the GOE (*13, 14, 21*). Somewhere in the range of 95 to 214 million years then passed (depending on where in the lineage each gene evolved, Figure 3, Table 1) before further genes involved in septal maturation (such as *hglK*) and regulation of cellular differentiation (namely *hetZ* and *patU3*) evolved at the start of the GOE in the most recent common ancestors of Macro- and Micro-cyanobacteria (Figure 3).

Crucial for the proliferation of primary producers was the emergence of evolutionary adaptations that enabled the formation of robust laminated microbial mats (*12*). These likely facilitated ecological expansions via shielding from lethal (ultraviolet) radiation and improved attachment to new substrates, resulting in oxygen production on a scale large enough to alter Earth’s atmosphere (*14*). Various factors contribute to the ability to form microbial mats, including large cell diameters, the presence of sheaths and motility (*12, 64*). One of these features, large cell size, is characteristic of Macrocyanobacteria and the Crown group of this clade emerged shortly after the evolution of genes (namely *hglK, patU3* and *hetZ*) guiding septal maturation and cellular differentiation in filaments at the start of the GOE ∼2.45 Ga (Figure 3, Node 6, Table 1). Therefore, we find that the timing of the emergence of multicellularity genes followed by the emergence of Macrocyanobacteria coincides broadly with atmospheric oxygenation and geological records documenting mats with increased tensile strength and cohesion in the late Archean (*73*).

### Intercellular Communication is not Limited to Heterocystous Strains

Some filamentous cyanobacteria that lack heterocysts (morphological section III, e.g. *Oscillatoria terebriformis*) do not exchange metabolites between cells in the laboratory (*44*), but others do. For example, *Microcoleus vaginatus*, a non-heterocystous species (*74*), has been found to exhibit intercellular communication at rates similar to those of *Anabaena* (*75*). Here, we have demonstrated that four further non-heterocystous strains from diverse positions in the cyanobacterial tree of life (including early-branching strains, Micro- and Macro-cyanobacteria) can also exchange substances between cells along their filament, despite the absence of heterocysts (Table 3). Additionally, they show regulation of septal junction function since both communicating and non-communicating cells are observed in all cases. These strains diversified from one another ∼ 2.6 Ga (Figure 3, node 3, Table 1), so it is possible that this trait has been a feature of filamentous cyanobacteria since their evolution in the Neoarchean (Figure 3, Table 1).

Whether intercellular communication was as fast in these early filamentous lineages as in Nostocales remains uncertain because the Micro- and Macro-cyanobacteria studied here show in general slower intercellular transfer of fluorescent markers than *Anabaena*, and the number of nanopores per septal disk is also generally smaller than in *Anabaena* (Table 3). This may be due to subtle differences in the Micro- and Macro-cyanobacterial septal proteins SepJ, FraC and FraD because, for example, point mutations in *Anabaena* SepJ have been shown to affect calcein transfer (*46*).

*Pseudanabaena* spp. are different to other filamentous cyanobacteria because they lack many of the septal junction proteins and cell differentiation genes found in filamentous Macro- and Micro- cyanobacteria, including the Nostocales (Figure 1). Perhaps as a result, *Pseudanabaena* sp. PCC 7367 has a unique septal disk perforated by a single pore instead of the more numerous smaller pores found in filamentous Macro- and Micro-cyanobacteria and *Anabaena* (Table 3). A similar single central pore has also been observed in *Oscillatoria lutea*, but it is smaller encompassing 23.32 ± 0.52 nm (*75*) instead of 40.10 ± 20 nm in *Pseudanabaena* sp. PCC 7367 (Table 3). Neither strain has been found to transfer calcein along their filaments, but we have shown that *Pseudanabaena* sp. PCC 7367 can transfer 5-CF which is significantly smaller than calcein (*69*).

*Pseudanabaena* sp. PCC 7367 appears to lack the FraD-containing septal junctions involved in transfer of calcein and 5-CF, so it must contain another type of junction specifically transferring 5-CF and metabolites of a similar size (about 376 Da). Whether these hypothetical junctions require SepJ to be formed remains to be determined, but it has been observed that filaments of *Pseudanabaena* spp. are generally short, consisting of less than 3 to 10 cells (*18*). This likely prevents them from establishing thick laminated microbial mats, which have not been observed for this clade (*18*). However, as the lineage leading to *Pseudanabaena* spp. emerged ∼ 2.6 Ga (node 3; Table 1, Figure 3, 95% credibility intervals span 2.80 to 2.38 Ga), around the same time as early ‘whiffs’ of oxygen appeared in shallow-ocean environments between ∼ 2.7 and ∼ 2.5 Ga in Western Australia and South Africa (*76-78*), we propose that *Pseudanabaena-*like ancestors represent earlier evolutionary steps contributing to biological changes in the lead up to the GOE (*58, 59, 65*). More specifically, we argue that early filamentous communities were responsible for early ‘whiffs’ of oxygen in marine environments, but that in addition to filament formation, other traits were also needed (e.g., sheaths, pigments, large cells, nitrogen fixation and UV-absorbing pigments such as scytonemin) to facilitate the rapid diversification of ancestral cyanobacteria into different habitats across land and marine coastal environments (*12, 79, 80*).

The purpose of intercellular communication in non-heterocystous strains remains unknown, but may help to synchronize circadian rhythms throughout the filaments. Under N-sufficient conditions, in which *Anabaena* filaments are composed only of vegetative cells, intercellular communication enables each cell to synchronize their behaviour so that circadian gene expression remains spatially-coherent along each filament (*81*). This could be a general property of filamentous cyanobacteria regardless of their evolutionary history, as it may enable all cells in the filament to function together as a single organismal unit.

### The Geological Record

Geochemical (*7, 10*) and biological evidence (*82*) point to an early origin of oxygenic photosynthesis in the early Archean. However, some major events in the early evolution of the biosphere continue to be controversial (*8*) due to uncertainties accompanying the interpretation of deep time events. Only a handful of studies have investigated the biological drivers that contributed to early diversification of Cyanobacteria and their subsequent ecological dominance across different habitats (*12*), so little is known about the biological controls that influenced physical Earth processes in the lead up to the planet’s oxygenation (*8*). Evolutionary studies addressing step changes in key primary producers like Cyanobacteria (e.g. (*59*) and (*83*)) are therefore needed to interpret geological events. Here, our molecular clock estimates of the last common ancestor (LCA) of Cyanobacteria ∼2.95 Ga (node 1, Figure 3, Table 1, 95% credibility intervals span 3.05 to 2.77 Ga) highlight the lag between the origin of the Cyanobacteria crown group and widespread geochemical evidence of oxygen at ∼ 2.4 Ga, in-line with several previous studies (e.g. (*1, 2, 21, 80, 84*)). Many geochemical hypotheses have been put forward to explain this, including a lack of phosphorus availability (*85, 86*) and competition for light and nutrients with photoferrotrophs (*87*).

### Evolution of branching filaments

Few cyanobacteria can grow branching filaments, and those that do belong to morphological section V. Our phylogenomic analyses reveal that strains from section V are polyphyletic (Figure 1), implying that the ability to grow branching filaments either evolved once and was subsequently lost in *Scytonema* spp., *Richelia* spp. and *Mastigocoleus testarum*, or it evolved independently as a result of convergent evolution. Previous studies have found that branches are created in *Mastigocladus laminosus* by randomized directions of cell growth (*45*). Perhaps alterations in cell division-related proteins determine changes in the angle of cell division. For example, inactivation of *mre* genes results in delocalization of the cell division ring in *Anabaena* (*88*).

### Conclusions

Our phylogenomic and molecular clock age estimates provide biological evidence unravelling the causes and consequences of perturbations of global carbon cycling and the rise in atmospheric oxygen in the Paleoproterozoic. By analysing the evolution of genes associated with the development of filaments in Cyanobacteria, we provide insight into evolutionary innovations which led to the GOE and one of life’s major transitions; the move from single isolated cells to interconnected cells in filamentous organisms. Phylogenetic reconstructions reveal that genes required to build septal junctions and create cell patterning are not restricted to heterocyst-formers but widespread amongst all filamentous Cyanobacteria, except for *Pseudanabaena* spp., which seem to have evolved a different or simpler method of growing filaments. Phylogenetic evidence suggests that some genes (including those encoding key septal proteins *sepJ* and *sepI*) were present before they diverged and have been inherited by all filamentous lineages, but others (including the cell differentiation genes *hetZ* and *patU3*) evolved later in the MRCA of filamentous Macro- and Micro-cyanobacteria. As a result, there were progressive increases in complexity, which began with *sepJ, sepI* and *fraE* in the ancestors of *Pseudanabaena*, and continued with the evolution of *hetZ* and *patU3* in the LCA of Macro- and Micro-cyanobacteria followed by the co-localisation of *sepJ* and *hetR* in the genomes of heterocystous strains. Future research efforts should focus on ascertaining the function of genes, such as those encoding the HetZ-related protein (encoded by *alr0202* in *Anabaena*) as they are good markers of filamentous morphology in Cyanobacteria, but their functions remain unknown.

## Materials and Methods

### Taxa Selection

Three novel cyanobacterial genomes were sequenced and a further 170 genomes downloaded from the NCBI (https://www.ncbi.nlm.nih.gov/). These taxa were chosen to embody the entire diversity of unicellular and filamentous cyanobacterial morphologies previously described by (*14, 19, 23, 60*) including all five morphological sections (Figure 1). Four genomes of Vampirovibrionia were included as outgroup (*5*).

Four strains are shown to illustrate the diversity of septal structures and the kinetics of intercellular communication in cyanobacteria from previously understudied groups (morphological section III): *Phormidesmis priestleyi* ULC 007, *Pseudanabaena* sp. PCC 9330 (here renamed *Nodosilinea), Nodosilinea* sp. LEGE 07298 and *Pseudanabaena* sp. PCC 7367. *Nodosilinea* sp. PCC 9330 was grown in BG11 medium with NaNO_3_ as the nitrogen source and citrate instead of ferric ammonium citrate used in the original recipe (*19*) and *Pseudanabaena* sp. PCC 7367 was grown in artificial seawater medium (*89, 90*), modified to lack NaSiO_3_ and vitamins, at 30 °C in the light (ca. 25 to 30 μmol photons m^-2^ s^-1^) in shaken (100 rpm) liquid cultures. *Phormidesmis priestleyi* ULC 007 was grown in BG11 medium (*19*) at 20°C and 20 μmol photons m^-2^ s^-1^ without agitation and *Nodosilinea* sp. LEGE 07298 was grown in DSMZ medium 1678 (BA+50+B+N/2) at 28°C, 30 μmol photons m^-2^ s^-1^ and constant agitation (110 rpm).

Other strains attempted include an additional early-branching strain (*Pseudanabaena* sp. PCC 6903), two Macrocyanobacteria (*Lyngbya* sp. PCC 8106 and *Phormidium laminosum* Gom OH-1-pC11) and one Microcyanobacterium (*Prochlorothrix hollandica* PCC 9006). For the Macrocyanobacteria (grown in the same conditions as Nodosilinea sp. PCC 9330 described above), isolation of clean murein sacculi failed for *Lyngbya* sp. PCC 8106 and no clear micrographs of septal disks nor any staining for FRAP analysis could be obtained from *Phormidium laminosum* Gom OH-1-pC11. For the Microcyanobacterium *Prochlorothrix hollandica* PCC 9006 (grown in BG11o + 2mM NaNO_3_ at 20⁰C without agitation and with 20 µmol photons m^-2^ s^-1^), not enough biomass for murein isolation was obtained. For the early branching strain, *Pseudanabaena* sp. PCC 6903 (grown in the same conditions as *Nodosilinea* sp. PCC 9330 described above), no clear micrographs of septal disks were obtained and FRAP analysis were difficult because of very small, motile filaments.

### Genome Sequencing and Assembly

Genomes from three novel filamentous strains from section III were sequenced. *Phormidium* sp. CCY1219 (synonymous to DSM 101373) and *Lyngbya* sp. CCY1209 (synonymous to DSM 101363) were obtained from Culture Collection Yerseke, having been originally isolated from sediments present in salt beds of Olhão, Portugal. Their mono-phototrophic cultures were isolated by plating and grown in ASN3 medium at 20⁰C on a 16h:8h light:dark cycle with 10-20 µmol m^-2^ s-^1^ of white light. Genomic DNA was extracted from 1.8 ml of culture with a DNeasy ultraclean microbial kit (Qiagen, Germany) according to the manufacturer’s instructions and stored in 10 mM Tris buffer at pH 8 and -80 ⁰C until paired-end reads of 2 x 150 bp were sequenced on a NextSeq 500/550 (Illumina, San Diego, CA) at the University of Bristol Genomics Facility, UK.

*Nodosilinea* sp. PCC 9330 was obtained from the Pasteur Culture Collection. It was grown in BG11 medium (see above) at 30 °C in the light (ca. 25 to 30 μmol photons m^-2^ s^-1^) in shaken (100 rpm) liquid cultures. Its genomic DNA was extracted from an axenic 50 ml of culture using the Molecular Biology Kit (Bio Basic) according to the manufacturer’s instructions. The cells were harvested by centrifugation (3000 x g, 5 min), resuspended in 50 ml of distilled water and the protocol was modified to include 0.1 mm glass beads (Qiagen, Germany) to increase DNA extraction by mechanical lysis. Purified genomic DNA from this culture was stored in nuclease-free water at 4 °C until paired-end reads of 2 x 250 bp were sequenced on an Illumina NextSeq500 at the Centro Andaluz de Biología Molecular y Medicina Regenerativa, Seville, Spain.

All reads were trimmed and assembled into draft genomes using the methods described in (*91*). Briefly, this included trimming sequences of poor quality base pairs with Trimmomatic v. 0.39 (*92*) and assembling the resulting reads into contigs using SPAdes v3.14.0 (*93*). Any non-cyanobacterial contigs were identified using BLAST and removed using a de bruijn graph visualisation approach developed for the de novo assembly of *Phormidesmis priestleyi* BC1401 (*94*). All draft genomes were submitted to JGI IMG/ER (*95*) for annotation (GOLD Analysis Project IDs: Ga0436897, Ga0436900 and Ga0591443) and deposited in DDBJ/ENA/Genbank repositories within BioProject PRJNA1053249 with BioSample accessions SAMN38850749, SAMN38850750 and SAMN38850751. Assembly statistics are presented in Table S3.

### Phylogenomic Tree of Cyanobacteria

To generate the phylogenomic tree of Cyanobacteria a total of 139 orthologous proteins and two ribosomal 23S LSU rRNA and 16S SSU rRNA were analysed; these genes are universally present in cyanobacterial taxa, evolutionarily conserved and had a minimum number of gene duplications (*1, 12, 21, 58, 96*). Genes were collected using the basic local alignment search tool for proteins (BLASTP) or nucleotides (BLASTN) with an e-value cut-off ≤ 1 x 10^-25^ and nine query sequences from diverse cyanobacteria (Table S4).

Homologs of these proteins and rRNA sequences were aligned with MAFFT v7.427 (*97*) and trimmed to remove gaps present in ≥ 85% of sequences using AlignmentViewer (available at http://sdsssdfd.altervista.org/arklumpus/AlignmentViewer/AlignmentViewer.html). Abnormally short and/or phylogenetically unique sequences (totalling 337 out of 23,961, just 1.41%) were manually removed. The best substitution model for each protein and rRNA was then chosen using ModelFinder (*98*). Maximum Likelihood was implemented to estimate the cyanobacterial tree of life in IQ-TREE v1.6.7 (*99*) using the -sp option to account for heterotachy (described in (*100*)). Support for branching relationships was estimated by calculating 1000 replicates of ultrafast bootstrap analyses (*101*) and the SH-like approximate likelihood ratio test (*102*). The topology was rooted using an outgroup consisting of four Vampirovibrionia, which are known to be the sister Phylum of Cyanobacteria (*5*). Trees were visualised in TreeViewer v. 2.2.0 (*103*).

### Distribution of Genes Involved in Septal Junction Formation

To investigate how strains from morphological section III generate and maintain their filaments, we searched for homologs of eight genes known to be involved in septal junction formation and the production of long filaments (Table S5). In the first instance, BLASTP searches were conducted to search for these proteins in the genomes used to generate our phylogenomic tree, using a permissive e-value cut-off ≤ 1 x 10^-5^ to ensure that no orthologs were missed. The resulting homologs were then aligned with MAFFT v7.427 (*97*) and used to construct ML trees in FastTree v2.1.10 (*104*). Any phylogenetically distant proteins were removed to ensure that all the remaining homologs were closely related to the query sequences.

Further stringency criteria were applied to remove homologs of HglK, SepJ and SepI with unrelated functions. Previous research has revealed that HglK contains a pentapeptide repeat region of 192 amino acids which is present in other proteins with different functions (*41, 105*). Whereas the active protein required for *Anabaena* to generate long filaments contains 727 amino acids (*41*), here we found that alignment lengths of HglK homologs vary from 767 to 41 amino acids with a noticeable drop-off below 700 (Figure S11), so only hits with alignment lengths > 700 were retained.

Similarly, SepJ has a domain which is closely-related to permeases from the drug/metabolite transporter family (DMT). These have no known role in filament formation, but are present in some filamentous taxa (*20, 43, 48*). Unlike SepJ, DMT permeases lack N-terminal coiled-coils and an intermediate “linker” domain (*106*), so only proteins with N-terminal coiled-coils and C-terminal TMHs, as predicted by ncoils (*107*) and TMHMM v2.0 (*108, 109*) were considered SepJ.

Finally, SepI is characterised by the presence of N-terminal coiled-coils and a domain similar to the linker domain of SepJ (*110*). To prevent any spurious proteins from influencing our results, homologs lacking essential features were discarded.

### Phylogenetic Reconstruction

A Bayesian approach was implemented to study the evolution of eight proteins involved in filament development. First, proteins were aligned in MAFFT v7.427 (*97*), then gaps trimmed if present in ≥ 85% of sequences. Phylogenies were estimated in MrBayes v3.2.7a (*111*) with a mixed amino acid substitution model prior, invariant sites and gamma distributed rates or Phylobayes v4.1 (*112*) using the GTR+CAT+G substitution model. Convergence was tested by comparing at least 2 replicate independent chains in one of two ways, depending on the method of phylogenetic reconstruction. Trees reconstructed in MrBayes were assessed in Tracer v1.6 (*113*) and considered converged when their average standard deviation of split frequencies (ASDSF) were ≤ 0.01 and potential scale reduction factors (PSRFs) between 1.00 and 1.02. For trees reconstructed in Phylobayes, convergence was reached when all effective sample size and relative difference (assessed with Tracecomp implemented in Phylobayes) were > 50 and < 0.3, respectively, after the first 25 % of iterations were discarded as burn-in. The resulting phylogenies were rooted on nodes with low ancestor deviation values predicted by MAD v2.2 (*114*)(Table S6).

### Chronology of Proteins

To predict when proteins associated with filamentous growth first appeared in cyanobacteria, we assessed the incongruence between our protein phylogenies and species tree (see similar approaches (*1, 58, 115-119*)). Congruence between our phylogenomic tree of cyanobacteria and protein phylogenies were taken to indicate shared evolutionary history caused by vertical inheritance from the clade’s MRCA.

### Fossil Calibrations

A total of six calibration points were implemented across all Cyanobacteria, all of which have been applied previously (*1, 58, 96*). Briefly, cyanobacteria with akinetes were constrained to arise 1.6 to 1.888 Ga and cyanobacteria with apical cells between 1.7 and 1.888 Ga based on fossil evidence (*120-123*). In addition, endosymbionts were constrained to arise before their hosts, specifically *Richelia intracellularis* HM01 and *R. intracellularis* HH01 before *Hemiaulus* fossils formed 110 Ma (*124, 125*) and UCYNA before *Braarudosphaera begelowii* appeared 91 Ma (*126, 127*). Geochemical evidence of the Great Oxygenation Event is applied such that cyanobacteria first diversify prior to 2.32 Ga (*128*), but after molybdenum isotopes document ‘whiffs’ of atmospheric oxygen 3 Ga (*76*). Cyanobacteria were constrained to diversify from their sister phyla, Vampirovibrionia (*5*), after the late heavy bombardment finished 3.9 Ga (*129, 130*) using a root prior of mean 3,060 and standard deviation 404 Ma. Soft bounds were applied throughout to allow 5% of the prior probability density of all calibrated nodes to fall outside of the specified bounds.

Previous studies (e.g.(*1, 21, 58, 96, 131*)) have applied a sixth calibration to constrain origin of filamentous cyanobacteria to ≥ 1.9 Ga based on filamentous microfossils found in the Belcher Islands (*132*). We remove this calibration point to avoid circular arguments and test when filamentous cyanobacteria first evolved without imposing a constraint. Our results (presented in detail below) suggest that many of the genes underlying filament development emerged in the MRCA of *Pseudanabaena* and heterocyst-forming strains, so we ran a further analysis where we apply the filamentous microfossil calibration to this node to see how it impacts estimated divergence times. Together, these strategies encompass a range of views on where filamentous morphology arose in the tree of life of cyanobacteria.

### Divergence Time Estimation

Divergence times were estimated by implementing a Bayesian relaxed molecular clock approach in Phylobayes 4.1 (*112*) using the topology generated above (Figure 1). We applied the CAT-GTR substitution model and uncorrelated gamma multipliers (*133*) model on 16S SSU rRNA and 23S LSU rRNA in Phylobayes 4.1 (*112*). This model has been found to be most appropriate for cyanobacterial phylogenies (*59*). For all noncalibrated nodes, we used a birth–death prior on divergence times. All calibrations had soft bounds, allowing 0.05% of the prior density to fall outside the minimum–maximum interval of each calibration. Convergence of these molecular clock analyses was assessed with tracecomp and consensus trees generated with readdiv (both implemented in Phylobayes). Clocks were considered converged when all effective sizes surpassed 50 and all relative differences were less than 0.3after the first 25% of iterations were discarded as burn-in.

### Peptidoglycan Sacculi Isolation and Visualization

Filaments of *Phormidesmis priestleyi* ULC 007, *Nodosilinea* sp. LEGE 07298, *Nodosilinea* sp. PCC 9330 and *Pseudanabaena* sp. PCC 7367 were harvested by centrifugation (3000 x g, 5 min, room temperature). Cells of *Phormidesmis priestleyi* ULC 007 and *Nodosilinea* sp. LEGE 07298 were frozen in liquid nitrogen, stored at -80 °C and shipped on dry ice to Seville for further analysis. Peptidoglycan (murein) sacculi were then isolated by a method modified from that of (*134*). The pellets were resuspended in 1 ml PBS buffer and homogenized by sonication using a sonicator bath. The broken filaments were added dropwise to boiling 6% (w/v) SDS and kept in this boiling condition for 2 hours, whilst carefully adding distilled water to avoid desiccation. Samples were incubated overnight at 37 °C with gentle stirring. Subsequently, four boiling steps were carried out: (i) 1 hour of boiling and ultracentrifugation (Beckmann 90 Ti rotor, 320,000 x *g*, 25°C, 35 min) and sample resuspension in 3 ml of 3% (w/v) SDS, (ii) 2 hours of boiling, centrifugation as above, and pellet resuspension in 2 ml distilled water and 0.2 ml of 6% (w/v) SDS, (iii) 2 hours of boiling, washing procedure by centrifugation, resuspension in 1.5 ml of 50 mM sodium phosphate (pH 6.8) and chymotrypsin or trypsin treatment (100 µg/ml) overnight at 37 °C. The incubation was terminated by adding 0.5 ml distilled water and 0.75 ml 6% (w/v) SDS. Finally, (iv) 2 hours of boiling, sedimentation by centrifugation as above, washing by distilled water several times to remove SDS by centrifugation (20,000 x *g*, 25°C, 5 min), resuspension in 100 µl of distilled water. The samples were stored at 4 °C. Samples of *Phormidesmis priestleyi* ULC 007 and *Nodosilinea* sp. LEGE 07298 were further washed twice with 1% (v/v) PBS and subsequently with distilled water. The purified sacculi were deposited on Formvar-carbon film-coated copper grids and stained with 1% (w/v) uranyl acetate. All the samples were examined with a Zeiss LIBRA 120 PLUS electron microscope at 120 kV (Servicio de Microscopía, Universidad de Sevilla, Seville, Spain).

### 5-CF and calcein staining

In this study, 5-carboxyfluorescein (5-CF) and calcein were loaded into the cytoplasm of *Phormidesmis priestleyi* ULC 007, *Nodosilinea* sp. PCC 9330, *Nodosilinea* sp. LEGE 07298,, and *Pseudanabaena* sp. PCC 7367 cells using a protocol modified from the one described previously for calcein by Mullineaux, Mariscal, Nenninger, Khanum, Herrero, Flores and Adams (*44*) or for 5-CF by Merino-Puerto, Schwarz, Maldener, Mariscal, Mullineaux, Herrero and Flores (*40*). In brief, the kinetics of intercellular communication were measured by staining cyanobacterial cells with 5-CF or calcein (*40, 44*) and visualising them by confocal fluorescence microscopy. After bleaching a single cell, we measured fluorescence recovery after photobleaching (FRAP). Subsequent imaging can reveal recovery of fluorescent 5-CF or calcein in the bleached cell due to simple diffusion from adjacent cells through septal junctions (*69, 135*).

The AM (acetoxymethyl) ester derivatives of calcein and 5-CF were dissolved in dimethyl sulfoxide (DMSO) to a concentration indicated below. For calcein staining of *Nodosilinea* sp. PCC 9330 and *Pseudanabaena* sp. PCC 7367, a total of 0.5 ml of cell culture was harvested by gentle centrifugation (3,000 x g, 5 min) washed three times in the respective growth medium as described above, resuspended in 0.5 ml fresh medium, and mixed with 20 μl of calcein AM solution (Thermo Fisher Scientific, 1 mg ml^-1^). The cells were incubated in dim light at 30°C for 90 min before harvesting and washing 3 times in fresh BG11 or artificial seawater medium, respectively. Calcein staining of *Nodosilinea* sp. LEGE 07298 and *Phormidesmis priestleyi* ULC 007 was performed as described above with the following modifications. After centrifugation, cells were resuspended in 1 ml of the respective growth media and 12 µl calcein AM (Cayman Chemical; 1 mg ml^-1^) was added. The suspension was incubated in darkness for 90 min at 28 °C or 20 °C, respectively and further treated as described above. For 5-CF staining of *Nodosilinea* sp. PCC 9330 and *Pseudanabaena* sp. PCC 7367, a total of 1 ml of the cell culture was harvested by gentle centrifugation (3,000 x g, 5 min), washed three times in the respective growth medium, resuspended in 1 ml fresh medium, and mixed with 12 μl of 5-CF AM solution (Thermo Fisher Scienfic, 5 mg ml^-1^). The cells were incubated in dim light at 30 °C for 30 min before harvesting and washing 3 times in fresh medium. 5-CF staining of *Nodosilinea* sp. LEGE 07298 and *Phormidesmis priestleyi* ULC 007 was performed as described above but using 5-CF AM from Thermo Fisher Scientific at a concentration of 1 mg ml^-1^ and incubating the cells in complete darkness for 60 min at 28 °C or 20 °C, respectively.

For both markers, filament suspensions were spotted onto BG11-agar (filaments were allowed to settle down by drying off excess liquid), and small agar blocks with labelled filaments were transferred to a custom-built temperature-controlled sample holder. The temperature was kept at the respective growth temperatures during the FRAP assays.

### FRAP analysis

For 5-CF in *Nodosilinea* sp. PCC 9330, filaments were imaged with a Leica TCS SP5 confocal laser-scanning microscope with a 488-nm line argon laser as the excitation source. Fluorescence emission was monitored by collection across windows of 500 to 541 nm with a 150-µm pinhole. For 5-CF and calcein in *Pseudanabaena* sp. PCC 7367, and calcein in *Nodosilinea* sp. PCC 9330, filaments were imaged with an Olympus Fluoview FV300 with a 488-nm line argon laser as the excitation source. Fluorescence emission was monitored by collection across windows of 500 to 527 nm with a 150-µm pinhole. After an initial image was recorded, bleaching was carried out by an automated FRAP routine as previously reported (*44*). Post-bleach images were taken in XY-mode approximately every two seconds for up to 1 to 2 min.

Fluorescence images of *Nodosilinea* sp. LEGE 07298 and *Phormidesmis priestleyi* ULC 007 stained with either calcein or 5-CF were acquired with an inverted confocal laser scanning microscope (CLSM; Leica SP8) using a HC PL APO x63/1.40 oil objective. Cells were excited at 488 nm and fluorescence emission collected from 500-540 nm. The pinhole was maximally opened to 600 μm, resulting in an optical section thickness of ∼4.9 μm. After the bleach images were recorded every second for 35 s (5-CF) or every two seconds for up to 1 min (calcein). Images were analysed with Leica LAS X (version 3.7.4) and FIJI (*136*).

For FRAP data analysis, kinetics of fluorescent marker transfer between cells located in the middle of filaments was quantified by determining the recovery rate constant R from the formula C_B_ = C_0_ + C_R_ (1 – e-^2*R*t^), where C_B_ is fluorescence in the bleached cell, C_0_ is fluorescence immediately after the bleach and tending towards (C_0_ + C_R_) after fluorescence recovery, t is time, and *R* is the recovery rate constant due to transfer of the marker from the neighbouring cells (*135*). Analysis were performed with Microsoft Excel and OriginPro 2019 (OriginLab) or GraphPad Prism.

## Acknowledgements

We thank Andrew Knoll and Kurt Konhauser for in-depth discussions on the early fossil record. Phylogenetic analyses were performed at the High Performance Computer facility (BlueCrystal Phase 4) at the University of Bristol. We would also like to acknowledge the assistance of the Core Facility BioSupraMol supported by the Deutsche Forschungsgemeinschaft (DFG) for confocal microscopy at FU Berlin.

Funding support for this work came from a University Royal Society Fellowship to P.S.-B, a University of Bristol Graduate Teaching Scholarship to J.S.B, grant number PID2020-118595GB-I00 from Agencia Estatal de Investigación del Ministerio de Ciencia e Innovación (Spain) and the European Regional Development Fund to E.F., grant number PY20_00058 from the Regional Government of Andalucía to E.F, a Regional Government of Andalucía research contract to M.N.M, and a DFG Emmy Noether project award (no. NU421/1) to DJN.

## Data Availability

Data is available in the supplementary materials and in the open science repository here: https://datadryad.org/stash/share/g0hGVGnBjb4XY1JRkwfG3IBz9nMDXpn-LD-GC_mgIO0 (doi:10.5061/dryad.t4b8gtj84). Also included here are newick formats of the protein phylogenies, species tree of Cyanobacteria and molecular clocks.

## Author Contributions

J.S.B, P.S-B and E.F. conceptualized the project. E.F., D.N. and M.N-V designed the experimental analyses. M.N-M, S.A., D.J.N and E.F. investigated intercellular communication and visualised septal morphology. J.S.B and P.S-B designed the phylogenetic analyses, whilst J.S.B performed the phylogenetic analyses. E.F. and P.S-B supervised the project. J.S.B, M.N-M and P.S-B wrote the original draft. All authors contributed to review and editing.

