## Supplementary material for "Evolution of Multicellularity Genes in the Lead Up to the Great Oxidation Event"

### Supplementary Information for Manuscript Entitled ‘Evolution of Multicellularity Genes in the Lead Up to the Great Oxidation Event’

Joanne S. Boden[1,4†], Mercedes Nieves-Morión[2†], Dennis J. Nürnberg[3,5], Sergio Aravelo[2], Enrique Flores[2], Patricia Sánchez-Baracaldo[4*]

[1] School of Earth and Environmental Sciences, University of St. Andrews, St. Andrews, Scotland, UK

[2] Instituto de Bioquímica Vegetal y Fotosíntesis, CSIC and Universidad de Sevilla, Seville, Spain.

[3] Institute for Experimental Physics, Freie Universität Berlin, Berlin, Germany

[4] School of Geographical Sciences, University of Bristol, Bristol, BS8 1SS, United Kingdom

[5] Dahlem Centre of Plant Sciences, Freie Universität Berlin, Berlin, Germany

[†] These authors contributed equally to this work.

Includes:

- Figures S1 to S12
- Tables S1 to S6

**Figure S1: Bayesian phylogeny of CyDiv homologs from cyanobacteria.** Numbers at the intersection between nodes and the start of collapsed clades represented by grey (multicellular strains) and yellow (unicellular strains) triangles represent posterior probability values. The phylogeny was constructed in MrBayes v3.2.6 (*1*) from an alignment of 259 amino acid positions.

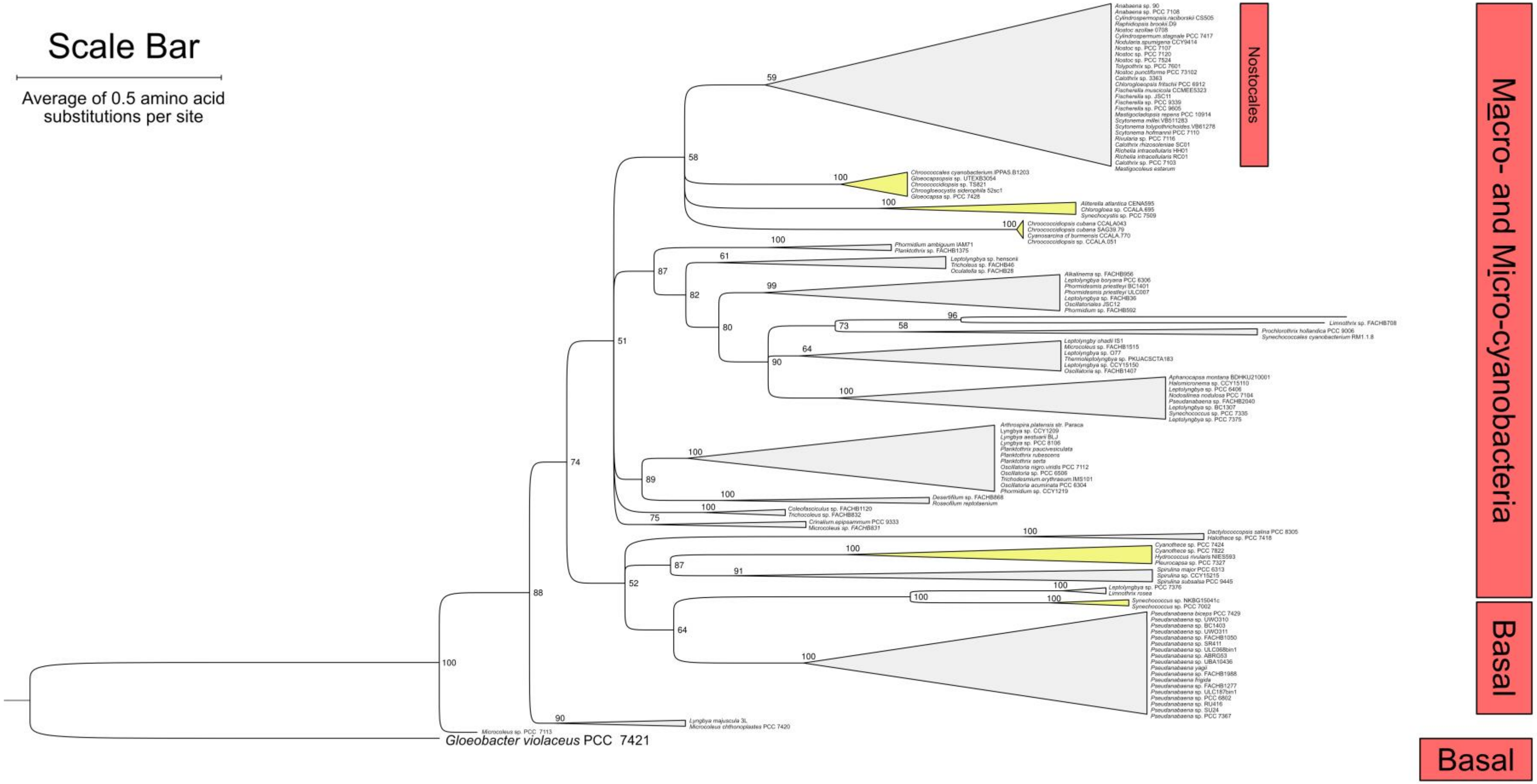

**Figure S2: Bayesian phylogeny or HetR homologs from cyanobacteria.** Numbers at the intersection between nodes and the start of collapsed clades represented by triangles are posterior probability values. Branches highlighted on a dark grey background represent secondary paralogs. The two HetR proteins of *Leptolyngbya* sp. FACHB261 are highlighted in blue and green text. The blue homolog is truncated, whereas the green protein is not. The phylogeny itself was constructed in MrBayes v3.2.6 (*1*) from an alignment of 306 amino acid positions. Micro. Microcyanobacteria

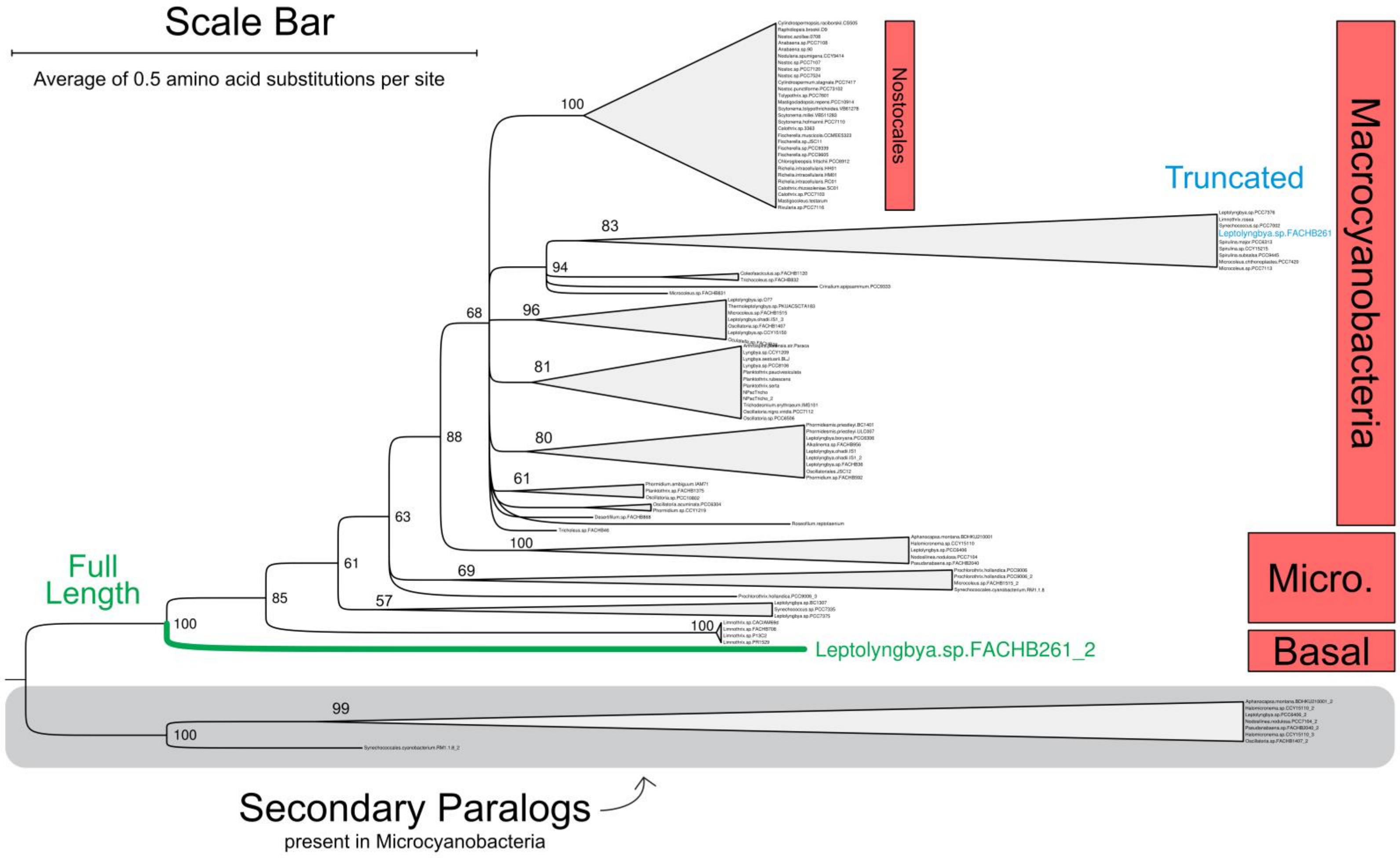

**Figure S3: Bayesian phylogeny of SepJ homologs from cyanobacteria.** Numbers at the intersection between nodes and the start of collapsed clades represented by grey triangles represent posterior probability values. The phylogeny was constructed in MrBayes v3.2.6 (*1*) from an alignment of 348 amino acid positions at the C terminus of the protein. Macro. Macrocyanobacteria, Micro. Microcyanobacteria.

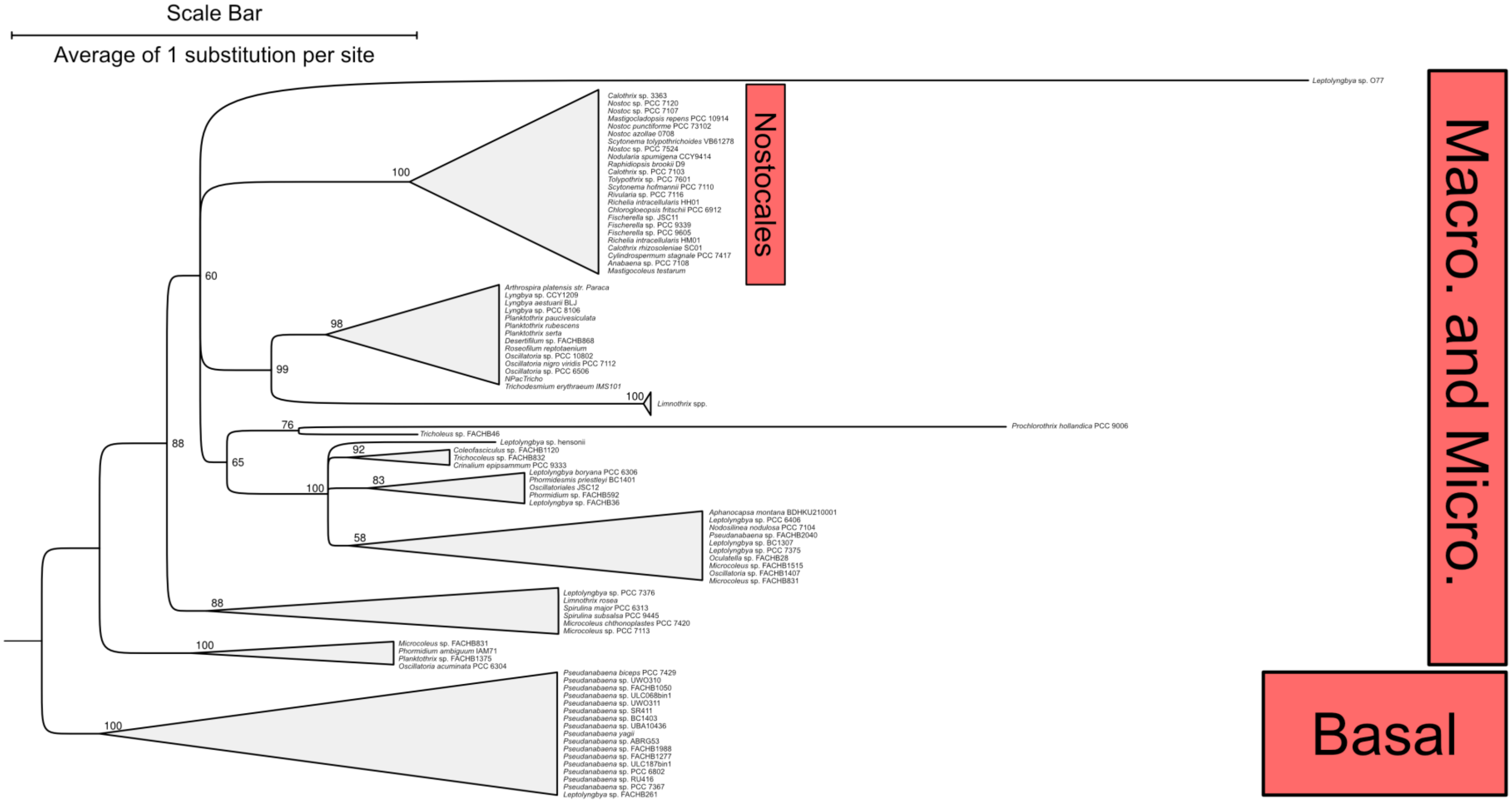

**Figure S4: Bayesian phylogeny of SepI homologs from cyanobacteria.** Numbers at the intersection between nodes and the start of collapsed clades represented by grey triangles represent posterior probability values. The phylogeny was constructed in MrBayes v3.2.6 (*1*) from an alignment of 195 amino acid positions including the coiled-coil region and linker domain at the C terminus. Micro. Microcyanobacteria.

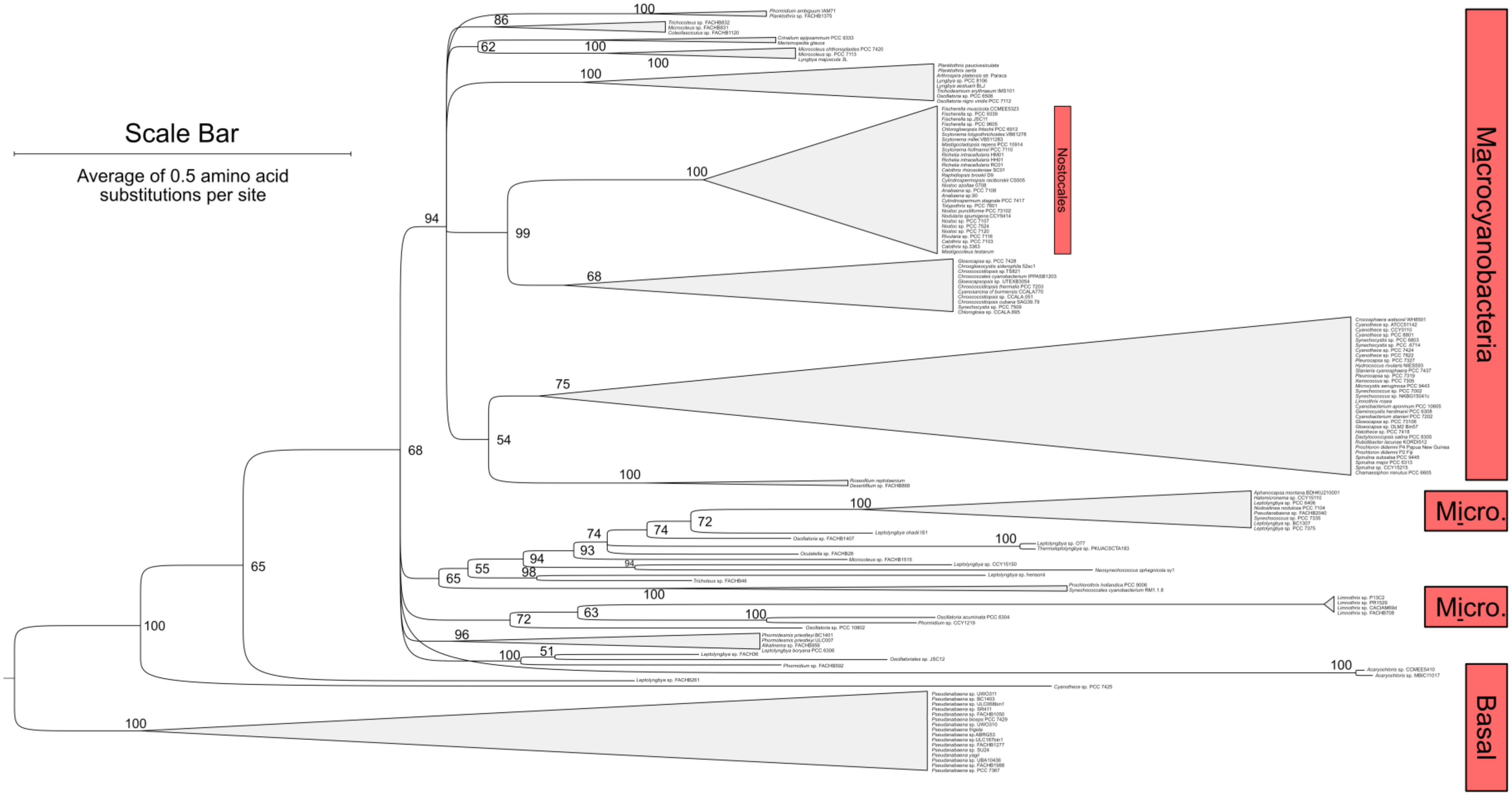

**Figure S5: Bayesian phylogeny of HetZ homologs (discounting 'HetZ related protein 2' for simplicity of viewing) from cyanobacteria.** The phylogeny is divided into two fields, here labelled as 'HetZ' and 'Related'. The field named 'HetZ' contains proteins named "heterocyst differentiation protein HetZ" on the NCBI (https://www.ncbi.nlm.nih.gov/) whereas the field named 'Related' contains proteins named "HetZ-related protein" on the NCBI (https://www.ncbi.nlm.nih.gov/). Numbers at the intersection between nodes and the start of collapsed clades represented by white triangles represent posterior probability values. The phylogeny was constructed in MrBayes v3.2.6 (*1*) from an alignment of 416 amino acid positions. Micro. Microcyanobacteria. Macro. Macrocyanobacteria.

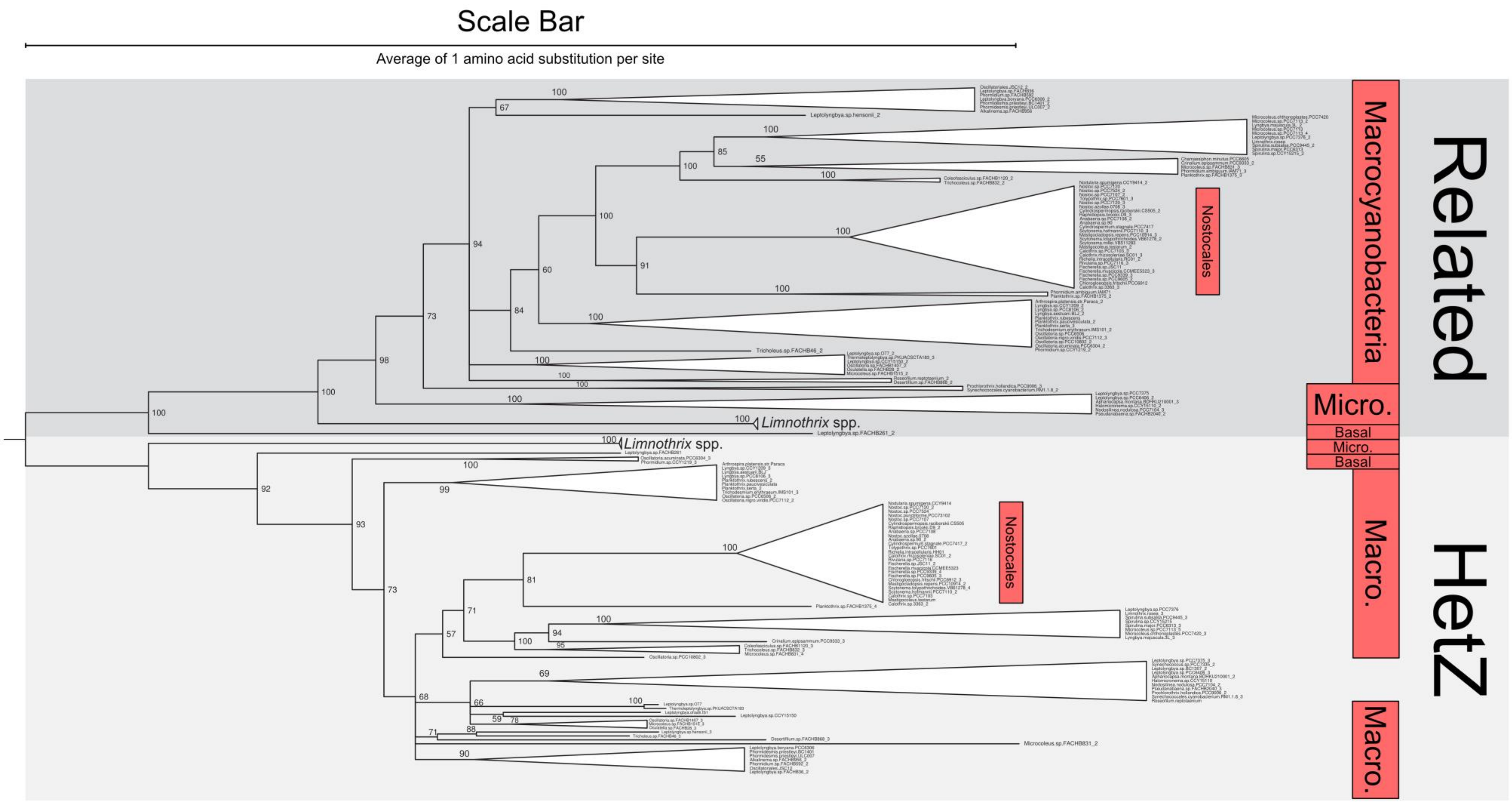

Scale Bar
Average of 1 amino acid substitution per site
Related
Macrocyanobacteria
Nostocales
Micro.
Basal
Micro.
Basal
Limnothrix spp.
Limnothrix spp.
HetZ
Macro.
Nostocales
Macro.

**Figure S6: Bayesian phylogeny of FraE homologs from cyanobacteria.** Numbers at the intersection between nodes and the start of collapsed clades represented by grey (multicellular strains) and yellow (unicellular strains) triangles represent posterior probability values. The phylogeny was constructed in MrBayes v3.2.6 (*1*) from an alignment of 264 amino acid positions. Micro. Microcyanobacteria.

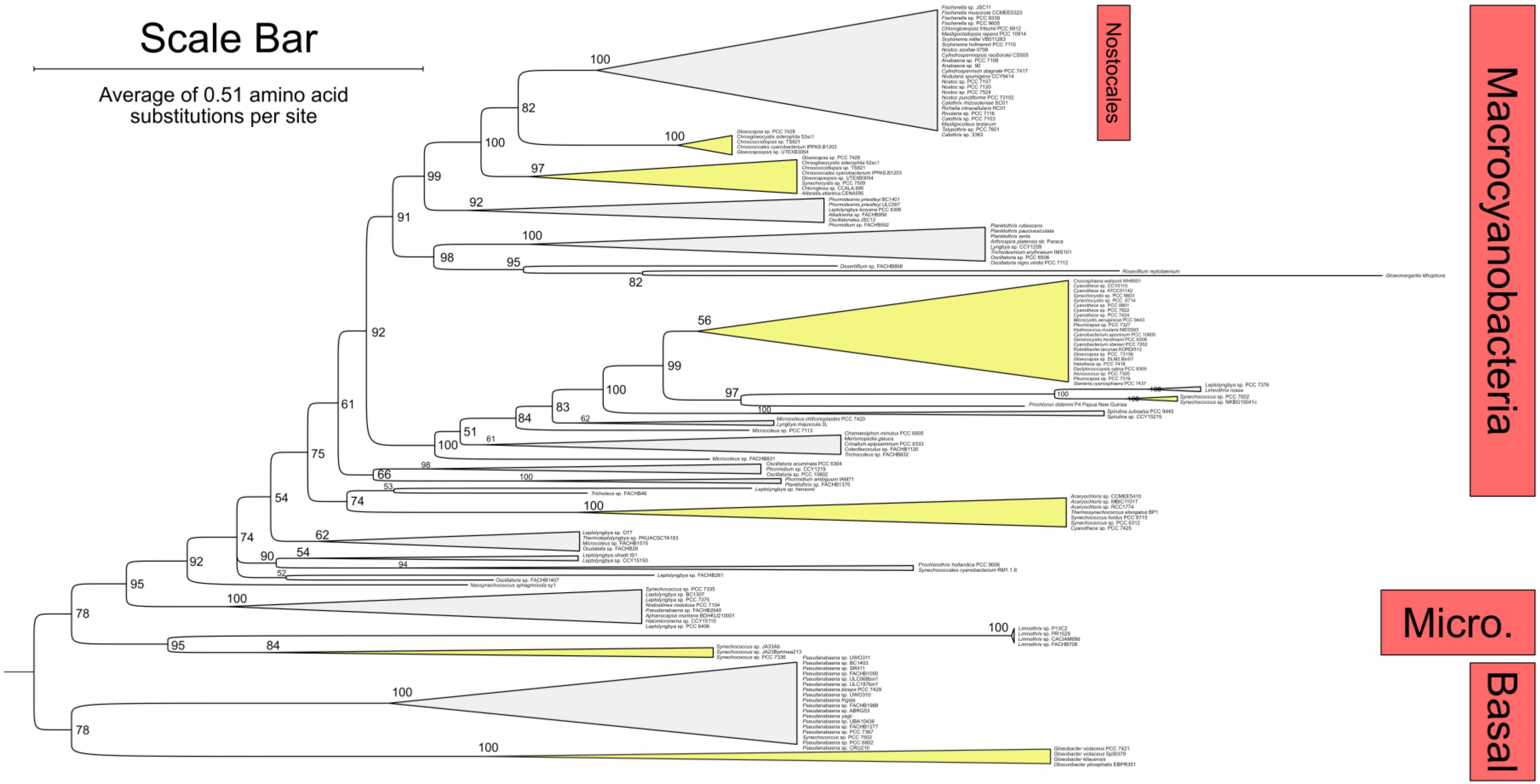

**Figure S7: Bayesian phylogeny of PatU3 homologs from cyanobacteria.** Numbers at the intersection between nodes and the start of collapsed clades represented by grey triangles represent posterior probability values. The phylogeny was constructed in MrBayes v3.2.6 (*1*) from an alignment of 374 amino acid positions. Micro. Microcyanobacteria

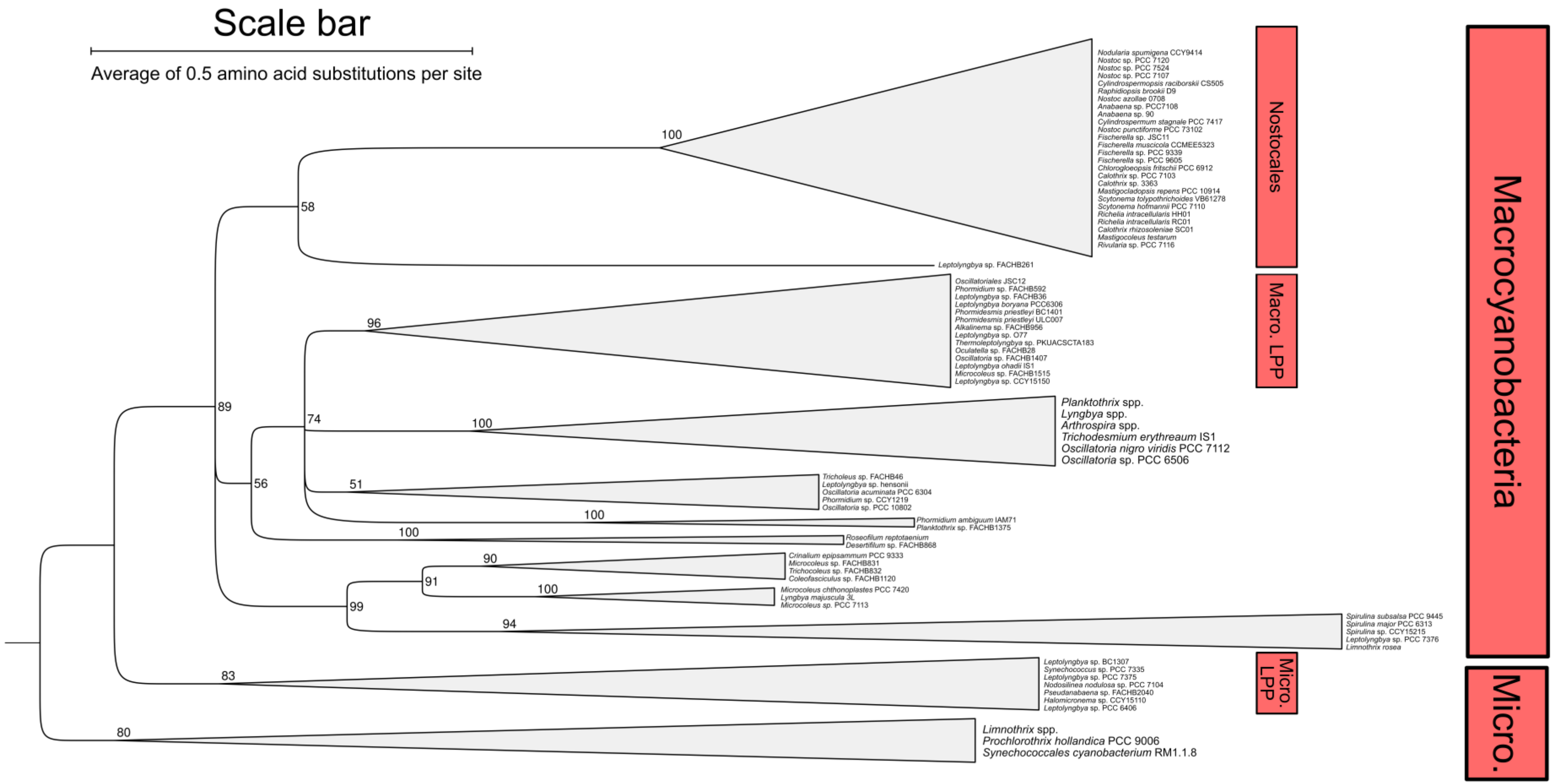

**Figure S8: Bayesian phylogeny of HglK homologs from cyanobacteria.** Numbers at the intersection between nodes and the start of collapsed clades represented by grey (multicellular strains) and yellow (unicellular strains) triangles represent posterior probability values. The phylogeny was constructed in Phylobayes v4.1 (*2*) from an alignment of 742 amino acid positions. Micro. Microcyanobacteria.

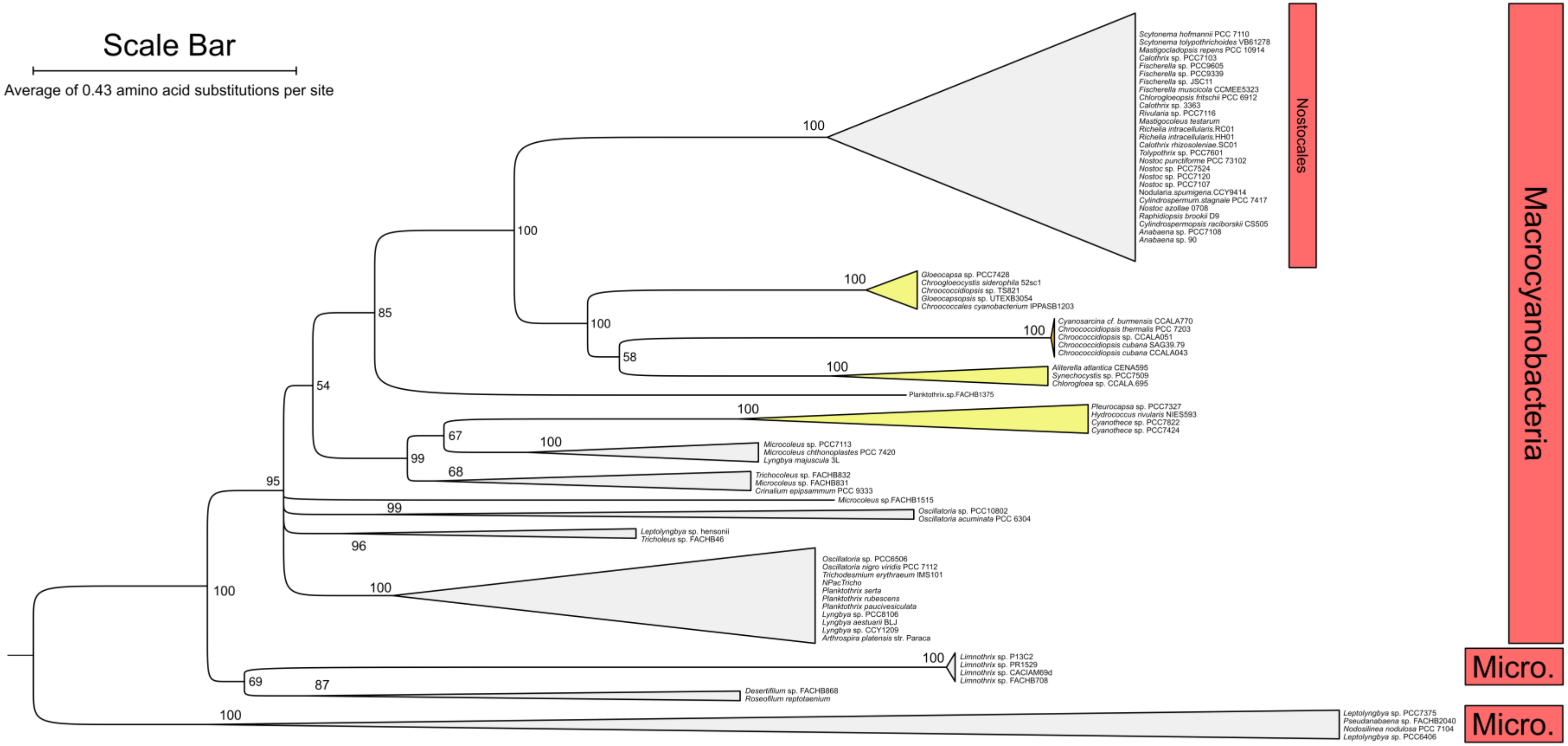

**Figure S9: Bayesian phylogeny of FraC homologs in cyanobacteria.** Numbers at the intersection between nodes and the start of collapsed clades represented by grey (multicellular strains) triangles represent posterior probability values. The scale bar in the top left signifies an average of 0.5 amino acid substitutions per site. The phylogeny was constructed in MrBayes v3.2.6 (*1*) from an alignment of 195 amino acid positions. Micro. Microcyanobacteria, Macro. Macrocyanobacteria.

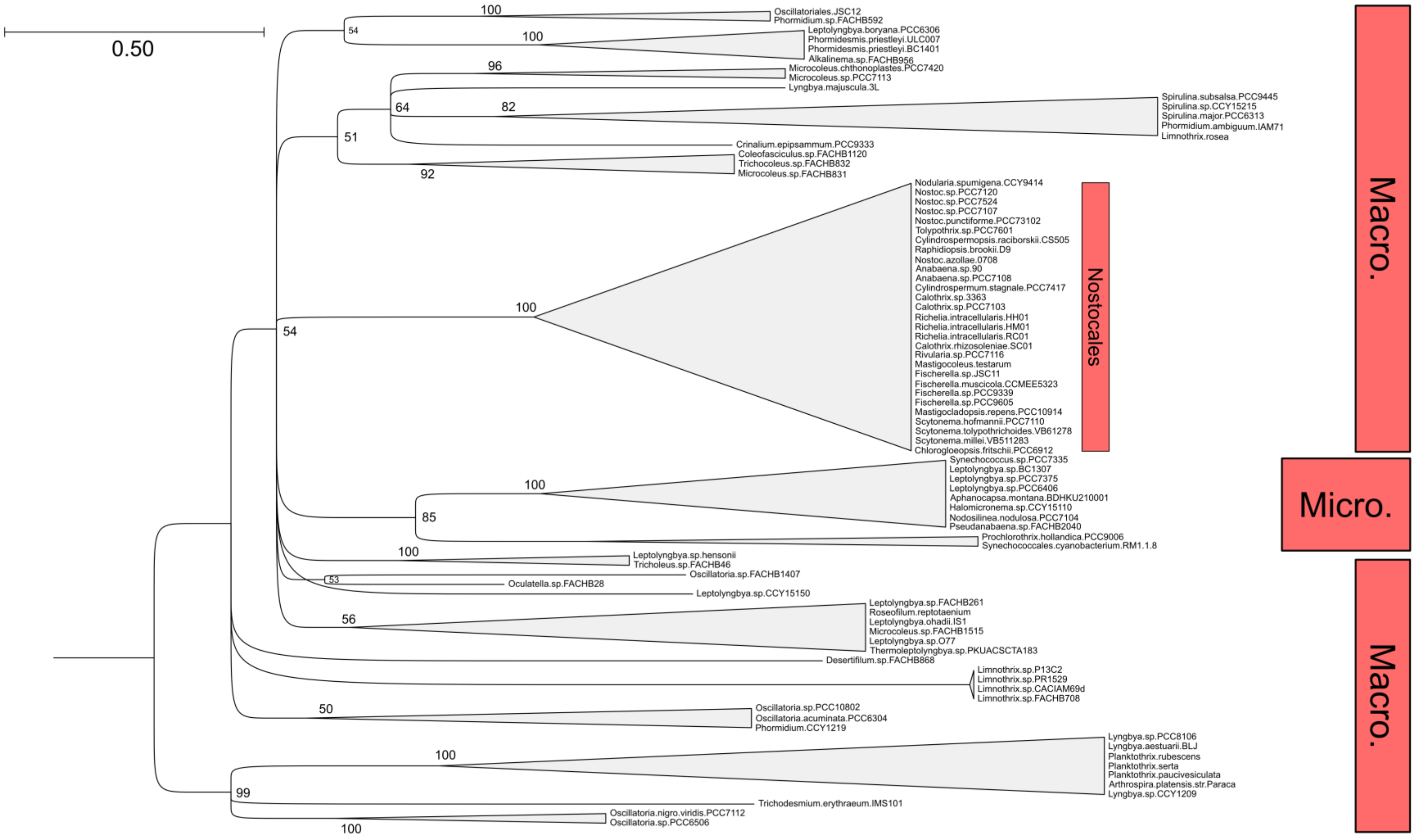

**Figure S10: Bayesian phylogeny of FraD homologs in cyanobacteria.** Numbers at the intersection between nodes and the start of collapsed clades represented by grey (multicellular strains) triangles represent posterior probability values. The phylogeny was constructed in MrBayes v3.2.6 (*1*) from an alignment of 356 amino acid positions. Micro. Microcyanobacteria, Macro. Macrocyanobacteria. Note that this rooting has the same minimal ancestor deviation (0.170) as an alternative root position (yellow star), which places sequences from a subset of Macrocyanobacteria (namely *Arthrospira* spp., *Planktothrix* spp. and some *Oscillatoria* spp.) as sisters to those of all other cyanobacteria.

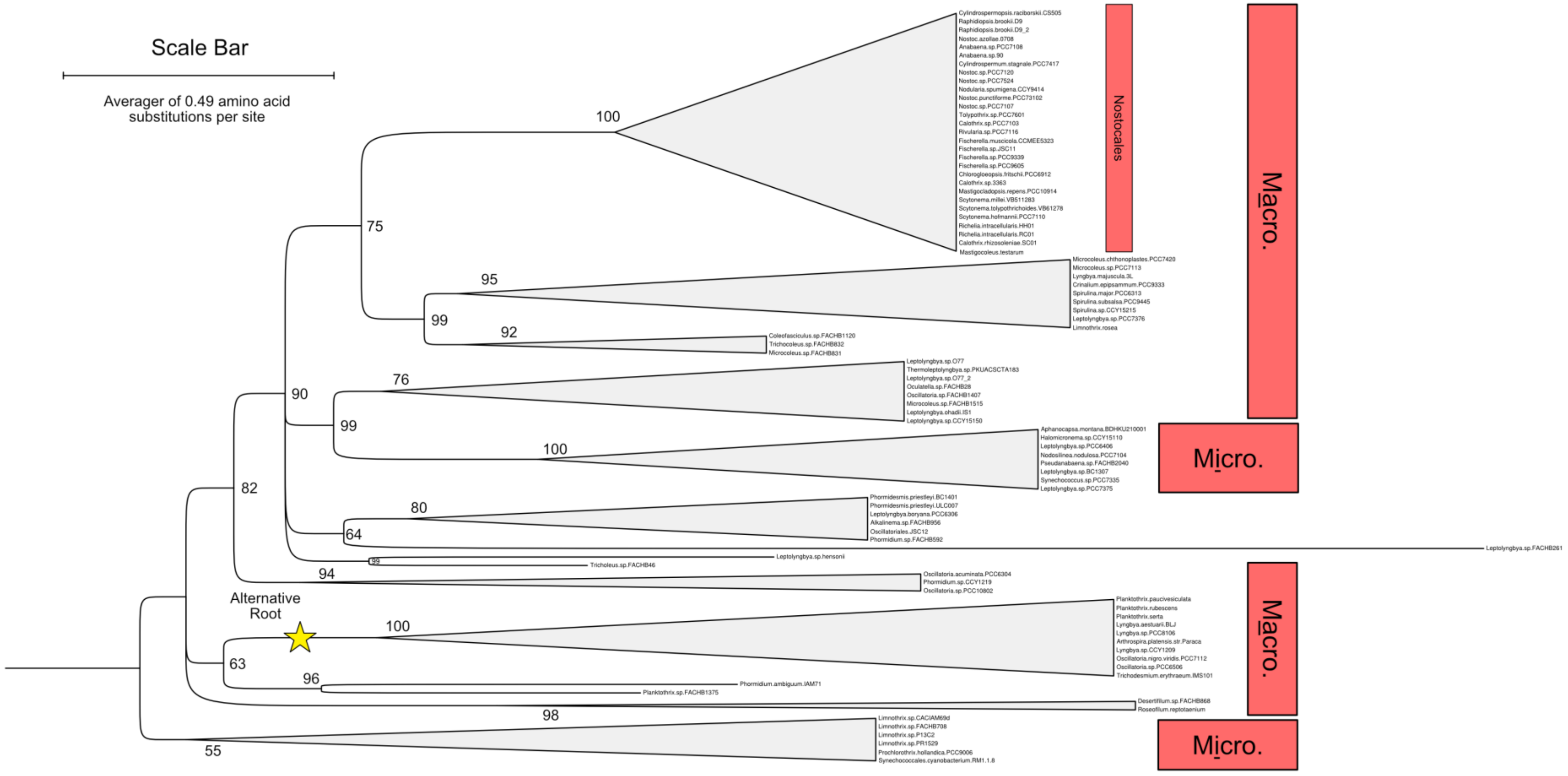

**Figure S11: Alignment length of HglK homologs in cyanobacteria.** Two y axis are used so that extremely frequent and less frequent homologs can be mapped on the same graph. Grey bars map to the left-hand axis because there are more than 1,000 homologs with the relevant alignment lengths. Whereas black bars map to the right-hand axis because there are less than 250 homologs with the relevant alignment lengths.

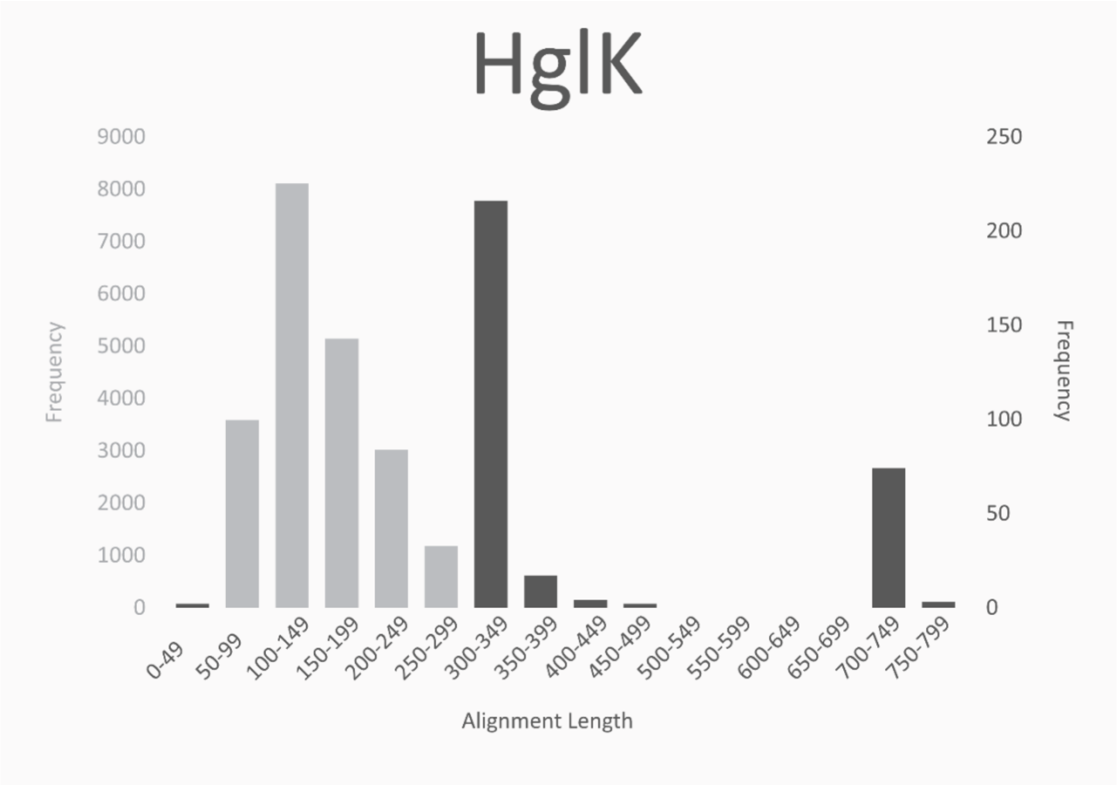

**Figure S12: Evolutionary trajectory of genes encoding septal proteins and components of cellular differentiation (pink, orange, blue and green lines) in the Cyanobacterial tree of life (black lines).** Interesting clades or strains of filamentous cyanobacteria are present at the tips of each branch (black text) above the genes typically found in that strain or clade (coloured squares matching the colour of the evolutionary trajectory of the gene it represents). Some genes have appeared in *Leptolyngbya* sp. FACHB261 following horizontal gene transfer from other cyanobacteria (dark grey cloud background), but most have been inherited vertically (plain white background) as shown by lines of matching colour on the evolutionary tree. The evolutionary history of a few genes (on light grey cloud backgrounds) remains unknown. All branch lengths are arbitrary.

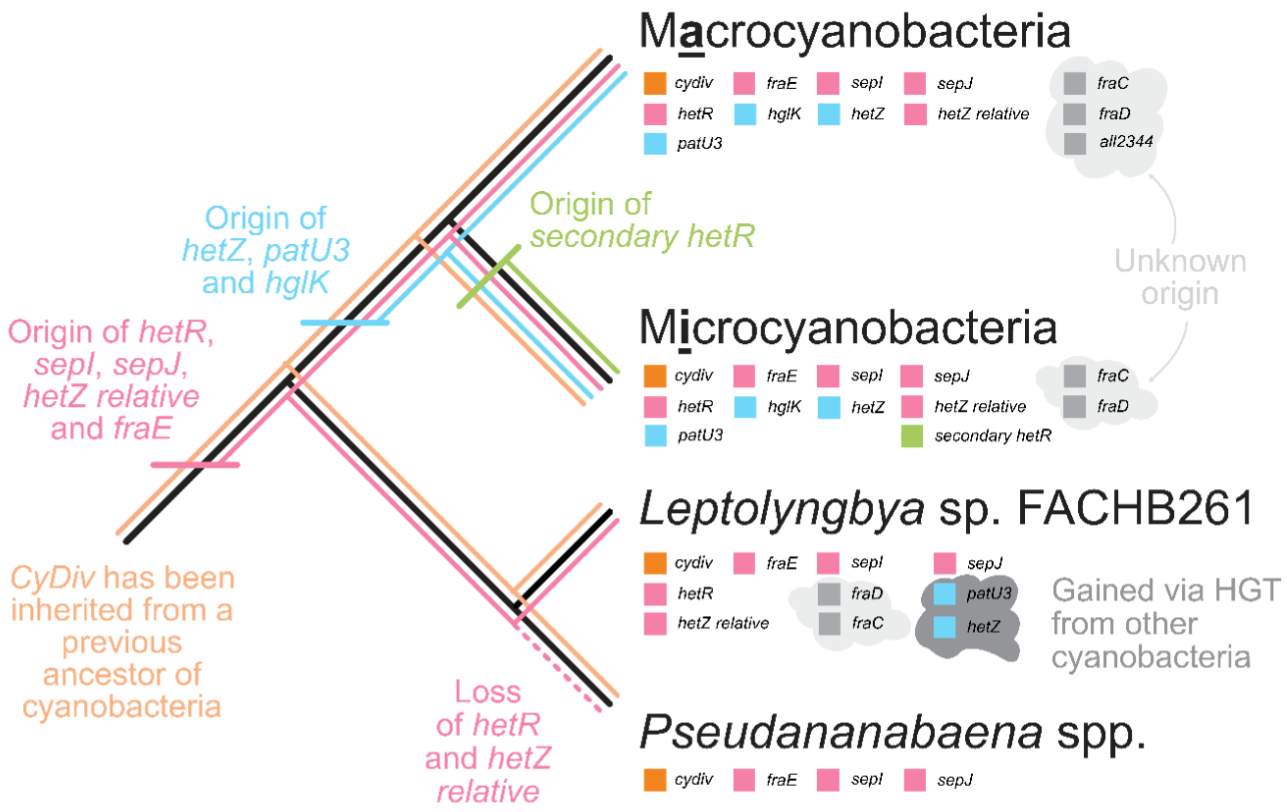

**Table S1: List of cyanobacteria and their morphological sections.** Colours applied to the morphological section column match those of Figures 1-3 to ease communicability.

| Strain | Morphological Section | Reference |
|---|---|---|
| Acaryochloris.sp.CCMEE5410 | I | (*3*) |
| Acaryochloris.sp.MBIC11017 | I | (*4*) |
| Aliterella.atlantica.CENA595 | II | (*3*) |
| Anabaena.sp.90 | IV | (*4*) |
| Anabaena.sp.PCC7108 | IV | (*4*) |
| Aphanocapsa.montana.BDHKU210001 | I | (*3*) |
| Arthrospira.platensis.str.Paraca | III | (*3*) |
| Calothrix.sp.3363 | IV | (*3*) |
| Calothrix.sp.PCC7I03 | IV | (*4*) |
| Chamaesiphon.minutus.PCC6605 | I | (*3*) |
| Chlorogloeopsis.fritschii.PCC69I2 | V | (*3*) |
| Chroococcidiopsis.cubana.SAG39.79 | II | (*4*) |
| Chroococcidiopsis.thermalis.PCC7203 | II | (*5*) |
| Crinalium.epipsammum.PCC9333 | III | (*3*) |
| Crocosphaera.watsonii.WH850I | I | (*4*) |
| Cyanobacterium.aponinum.PCC10605 | I | (*3*) |
| Cyanobacterium.stanieri.PCC7202 | I | (*3*) |
| Cyanothece.sp.ATCC51142 | I | (*4*) |
| Cyanothece.sp.CCY0110 | I | (*3*) |
| Cyanothece.sp.PCC7424 | I | (*4*) |
| Cyanothece.sp.PCC7425 | I | (*4*) |
| Cyanothece.sp.PCC7822 | I | (*4*) |
| Cyanothece.sp.PCC8801 | I | (*4*) |
| Cylindrospermopsis.raciborskii.CS505 | IV | (*3*) |
| Cylindrospermum.stagnale.PCC7417 | IV | (*4*) |

| | | |
|---|---|---|
| Dactylococcopsis.salina.PCC8305 | I | *(4)* |
| Fischerella.muscicola.CCMEE5323 | V | |
| Fischerella.sp.JSC11 | V | *(3)* |
| Fischerella.sp.PCC9339 | V | *(3)* |
| Fischerella.sp.PCC9605 | V | *(3)* |
| Geminocystis.herdmanii.PCC6308 | I | *(4)* |
| Gloeobacter.violaceus.PCC7421 | I | *(4)* |
| Gloeocapsa.sp.PCC.73106 | I | *(4)* |
| Gloeocapsa.sp.PCC7428 | I | *(3)* |
| Gloeomargarita.lithophora | I | *(6)* |
| Halothece.sp.PCC7418 | I | *(4)* |
| Leptolyngbya.boryana.PCC6306 | III | *(3)* |
| Leptolyngbya.sp.PCC6406 | III | *(3)* |
| Leptolyngbya.sp.PCC7375 | III | *(3)* |
| Leptolyngbya.sp.PCC7376 | III | *(3)* |
| Lyngbya.aestuarii.BLJ | III | *(3)* |
| Lyngbya.majuscula.3L | III | *(4)* |
| Mastigocladopsis.repens.PCC10914 | V | *(3)* |
| Mastigocoleus.testarum | V | *(3)* |
| Microcoleus.chthonoplastes.PCC7420 | III | *(4)* |
| Microcoleus.sp.PCC7113 | III | *(3)* |
| Microcystis.aeruginosa.PCC9443 | I | *(3)* |
| Nodosilinea.nodulosa.PCC7104 | III | *(3)* |
| Nodularia.spumigena.CCY9414 | IV | *(3)* |
| Nostoc.azollae.0708 | IV | *(4)* |
| Nostoc.punctiforme.PCC73102 | IV | *(4)* |
| Nostoc.sp.PCC7107 | IV | *(3)* |
| Nostoc.sp.PCC7120 | IV | *(4)* |
| Nostoc.sp.PCC7524 | IV | *(3)* |
| Oscillatoria.acuminata.PCC6304 | III | *(3)* |

| | | |
|---|---|---|
| Oscillatoria.nigro.viridis.PCC7112 | III | (*3*, *5*) |
| Oscillatoria.sp.PCC10802 | III | (*3*) |
| Oscillatoria.sp.PCC6506 | III | (*5*) |
| P.marinus.str.MIT9301 | I | (*4*) |
| P.marinus.str.MIT9303 | I | (*4*) |
| P.marinus.str.NATL1A | I | (*4*) |
| Phormidesmis.priestleyi.BC1401 | III | (*7*) |
| Planktothrix.rubescens | III | (*4*) |
| Pleurocapsa.sp.PCC7319 | II | (*4*) |
| Pleurocapsa.sp.PCC7327 | I | (*4*) |
| Prochlorothrix.hollandica.PCC9006 | III | (*4*) |
| Pseudanabaena.biceps.PCC7429 | III | (*3*) |
| Pseudanabaena.sp.PCC6802 | III | (*4*) |
| Pseudanabaena.sp.PCC7367 | III | (*4*) |
| Raphidiopsis.brookii.D9 | IV | (*3*) |
| Richelia.intracellularis.HH01 | IV | (*3*) |
| Rivularia.sp.PCC7116 | IV | (*3*) |
| Rubidibacter.lacunae.KORDI512 | I | (*4*) |
| Scytonema.millei.VB511283 | IV | (*3*) |
| Scytonema.tolypothrichoides.VB61278 | IV | (*3*) |
| Spirulina.major.PCC6313 | III | (*4*) |
| Spirulina.subsalsa.PCC9445 | III | (*3*) |
| Stanieria.cyanosphaera.PCC7437 | II | (*3*) |
| Synechococcus.elongatus.PCC7942 | I | (*4*) |
| Synechococcus.sp.CB0205 | I | (*5*) |
| Synechococcus.sp.CC9605 | I | (*4*) |
| Synechococcus.sp.JA23Bprimea213 | I | (*4*) |
| Synechococcus.sp.JA33Ab | I | (*4*) |
| Synechococcus.sp.NKBG15041c | I | (*3*) |
| Synechococcus.sp.PCC6312 | I | (*3*) |

| | | |
|---|---|---|
| Synechococcus.sp.PCC7002 | I | (*4*) |
| Synechococcus.sp.PCC7336 | I | (*5*) |
| Synechococcus.sp.PCC7502 | III | (*4*) |
| Synechococcus.sp.RCC307 | I | (*5*) |
| Synechococcus.sp.RS9916 | I | (*5*) |
| Synechococcus.sp.WH5701 | I | (*4*) |
| Synechococcus.sp.WH7805 | I | (*4*) |
| Synechocystis.sp.PCC.6714 | I | (*3*) |
| Synechocystis.sp.PCC6803 | I | (*4*) |
| Synechocystis.sp.PCC7509 | I | (*3*) |
| Thermosynechococcus.elongatus.BP1 | I | (*4*) |
| Tolypothrix.sp.PCC7601 | IV | (*3*) |
| Trichodesmium.erythraeum.IMS101 | III | (*4*) |
| UCYNA | I | (*4*) |
| Xenococcus.sp.PCC7305 | II | (*3*) |

**Table S2: Clade Definitions and Genetic Complements of *fraC, fraD, fraE, hetZ, patU3, hetR* and *sepJ*.** Each row represents a separate strain or clade. Genes are coloured to match Figure 1. Where cells representing two or more of *fraC*, *fraD* and *fraE* are coloured, the relevant genes are separated by < I,000 nucleotides in the genome. The same is true of *hetZ* and *patU3*. Where cells representing *hetR* and *sepJ* are coloured, the two genes are separated by less than 3,000 nucleotides in the genome. P. present but not close to another gene in the table. 'P' indicates that a gene is present in a separate operon. 'n/a' is stated when the genome of the strain in question has not been sequenced.

| Clade Name | Strain | *fraC* | *fraD* | *fraE* | *hetZ* | *patU3* | *hetR* | *sepJ* |
|---|---|---|---|---|---|---|---|---|
| Leptolyngbya sp. FACHB261 | Leptolyngbya sp. FACHB261 | | | | | | P | P |
| Pseudanabaena | Pseudanabaena sp. PCC7367 | | | | | | | |
| | Pseudanabaena sp. PCC6802 | | | | | | | |
| | Pseudanabaena sp. RU416 | | | | | | | |
| | Pseudanabaena sp. CRU210 | | | | | | | |
| | Pseudanabaena sp. SU24 | | | | | | | |
| | Synechococcus sp. PCC7502 | | | | | | | |
| | Pseudanabaena sp. FACHB1277 | | | | | | | |
| | Pseudanabaena sp. ULC187binI | | | | | | | |
| | Pseudanabaena sp. FACHB1988 | | | | | | | |
| | Pseudanabaena sp. PCC6903 | n/a | n/a | n/a | n/a | n/a | n/a | n/a |
| | Pseudanabaena sp. ABRG53 | | | | | | | |
| | Pseudanabaena sp. UBA10436 | | | | | | | |
| | Pseudanabaena yagii | | | | | | | |
| | Pseudanabaena biceps PCC7429 | | | | | | | |
| | Pseudanabaena sp. UWO310 | | | | | | | |
| | Pseudanabaena frigida | | | | | | | |
| | Pseudanabaena sp. FACHB1050 | | | | | | | |
| | Pseudanabaena sp. SR411 | | | | | | | |
| | Pseudanabaena sp. ULC068binI | | | | | | | |
| | Pseudanabaena sp. BC1403 | | | | | | | |
| | Pseudanabaena sp. UWO311 | | | | | | | |

| | | | | | | | | |
|---|---|---|---|---|---|---|---|---|
| Micro-LPP | Leptolyngbya sp. BC1307 | | | | | | P | P |
| | Pseudanabaena persicina SAG 80.79 | n/a | n/a | n/a | n/a | n/a | n/a | n/a |
| | Synechococcus sp. PCC7335 | | | | | | | |
| | Leptolyngbya sp. PCC7375 | | | | | | P | P |
| | Leptolyngbya sp. PCC6406 | | | | | | P | P |
| | Aphanocapsa montana BDHKU210001 | | | P | P | | P | P |
| | Halomicronema sp. CCY15110 | | | P | | | | |
| | Pseudanabaena sp. FACHB2040 | | | | | | P | P |
| | Nodosilinea.nodulosa PCC7104 | | | | | | P | P |
| | Pseudanabaena sp. PCC9330 | n/a | n/a | n/a | n/a | n/a | n/a | n/a |
| | Leptolyngbya sp. PCC9324 | n/a | n/a | n/a | n/a | n/a | n/a | n/a |
| Prochlorothrix | Prochlorothrix hollandica PCC9006 | | | P | | | P | P |
| | Synechococcales cyanobacterium RM1.1.8 | | | | | | | |
| Limnothrix | Limnothrix sp. CACIAM69d | | | | | | P | P |
| | Limnothrix sp. FACHB708 | | | | | | P | P |
| | Limnothrix sp. P13C2 | | | | | | P | P |
| | Limnothrix sp. PR1529 | | | | | | P | P |
| Macro-LPP | Tricholeus sp. FACHB46 | | | | | | P | P |
| | Leptolyngbya sp. hensonii | | | P | | | | |
| | Neosynechococcus sphagnicola.sy1 | | | P | | | | |
| | Alkalinema sp. FACHB956 | | | | | | | |
| | Leptolyngbya boryana PCC6306 | | | | | | P | P |
| | Phormidesmis priestleyi BC1401 | | | | | | P | P |
| | Phormidesmis priestleyi ULC007 | | | | | | | |
| | Oscillatoriales JSC12 | | | | | | P | P |
| | Leptolyngbya sp. FACHB36 | | | | | | P | P |
| | Phormidium sp. FACHB592 | | | | | | P | P |
| | Oculatella sp. FACHB28 | | | | | | P | P |
| | Leptolyngbya sp. CCY15150 | P | | | | | | |

| | | | | | | | | |
|---|---|---|---|---|---|---|---|---|
| | Leptolyngbya ohadii IS1 | | | P | | | | |
| | Microcoleus sp. FACHB1515 | | | P | | | P | P |
| | Oscillatoria sp. FACHB1407 | | | P | | | P | P |
| | Leptolyngbya sp. O77 | P | | | | | P | P |
| | Phormidium laminosum Gom OH1pC11 | n/a | n/a | n/a | n/a | n/a | n/a | n/a |
| | Thermoleptolyngbya sp. PKUACSCTA183 | P | | | | | | |
| Desertifilum | Desertifilum sp. FACHB868 | | | | | | P | P |
| | Roseofilum reptotaenium | | P | | | | P | P |
| Oscillatoria | Oscillatoria sp. PCC10802 | | | | | | P | P |
| | Oscillatoria acuminata PCC6304 | | | | | | P | P |
| | Phormidium sp. CCY1219 | | | | | | | |
| | Oscillatoria nigro viridis PCC7112 | | | | | | P | P |
| | Oscillatoria sp. PCC6506 | | | | | | P | P |
| | NPacTricho | | | | | | P | P |
| | Trichodesmium erythraeum IMS101 | | | | | | P | P |
| | Arthrospira platensis str. Paraca | | | P | | | P | P |
| | Lyngbya sp. CCY1209 | | | | | | P | P |
| | Lyngbya.aestuarii.BLJ | | | | | | P | P |
| | Lyngbya sp. PCC8106 | P | P | | | | P | P |
| | Planktothrix.serta | | | | | | P | P |
| | Planktothrix.paucivesiculata | P | | | | | P | P |
| | Planktothrix.rubescens | | | | | | P | P |
| Trichocoleus | Microcoleus sp. FACHB831 | | | P | | | P | P |
| | Coleofasciculus sp. FACHB1120 | | | | | | P | P |
| | Trichocoleus sp. FACHB832 | | | | | | P | P |
| *Crinalium epipsammum PCC9333* | Crinalium epipsammum PCC9333 | | | | | | P | P |
| Phormidium | Phormidium ambiguum IAM71 | | | | | | P | P |
| | Planktothrix sp. FACHB1375 | | | | | | P | P |
| Microcoleus | Lyngbya majuscula.3L | | | P | | | | |

| | | | | | | | | |
|---|---|---|---|---|---|---|---|---|
| | Microcoleus chthonoplastes PCC7420 | | | | | | P | P |
| | Microcoleus sp. PCC7113 | | | | | | P | P |
| Spirulina | Spirulina subsalsa PCC9445 | | | P | | | P | P |
| | Spirulina major PCC6313 | | | | | | P | P |
| | Spirulina sp. CCY15215 | | | P | | | | |
| *Limnothrix rosea and Leptolyngbya* sp. PCC7376 | Leptolyngbya sp. PCC7376 | | P | P | | | P | P |
| | Limnothrix rosea | P | P | P | | | P | P |
| Nostocales | Calothrix sp. 3363 | | | | | | P | P |
| | Calothrix sp. PCC7103 | | | | | | | |
| | Mastigocoleus.testarum | | | | | | P | P |
| | Rivularia sp. PCC7116 | | | | | | | |
| | Calothrix rhizosoleniae SC01 | | | | | | P | P |
| | Richelia intracellularis RC01 | | | P | | P | | |
| | Richelia intracellularis HH01 | | | | | | P | P |
| | Richelia intracellularis HM01 | | | | | | P | P |
| | Scytonema hofmannii PCC7110 | | | | | | | |
| | Scytonema tolypothrichoides VB61278 | | | | | | | |
| | Mastigocladopsis repens PCC10914 | | | | | | | |
| | Scytonema millei VB511283 | | | | | | | |
| | Chlorogloeopsis fritschii PCC6912 | | | | | | | |
| | Fischerella sp. PCC9605 | | | | | | | |
| | Fischerella sp. PCC9339 | | | | | | | |
| | Fischerella muscicola CCMEE5323 | | | P | | | | |
| | Fischerella sp. JSC11 | | | | | | P | P |
| | Tolypothrix sp. PCC7601 | | | | | | | |
| | Nostoc sp. PCC7107 | | | | | | | |
| | Nostoc sp. PCC7120 | | | | | | | |
| | Nostoc sp. PCC7524 | | | | | | | |
| | Nostoc.punctiforme PCC73102 | | | | | | | |

| | | | | | | | |
|---|---|---|---|---|---|---|---|
| Nodularia spumigena CCY9414 | | | | | | | |
| Cylindrospermum.stagnale PCC7417 | | | | | | | |
| Anabaena sp. 90 | | | | | | | |
| Anabaena sp. PCC7108 | | | | | | | |
| Nostoc azollae 0708 | | | | | | | |
| Cylindrospermopsis raciborskii CS505 | | | | | | | |
| Raphidiopsis brookii D9 | | | | | | P | P |

**Table S3: Assembly statistics of draft genomes sequenced for this study.**

| | *Phormidium* sp. CCY1219 | *Lyngbya* sp. CCY1209 | *Nodosilinea* sp. PCC 9330 |
|---|---|---|---|
| Genome length (bp) | 7,535,552 | 6,681,614 | 5,788,900 |
| Genome coverage | 233 x | 130 x | 4,031 x |
| $N_{50}$ | 27,142 | 24,980 | 114,311 |
| Number of contigs | 977 | 716 | 120 |
| GC content | 50.63 % | 54.24 % | 57.11 % |
| Completeness* | 98.5 % | 99.1 % | 97.3 % |
| Protein-coding genes | 6,799 | 5,972 | 5,322 |
| Number of rRNAs | 3 | 4 | 2 |
| Number of tRNAs | 66 | 48 | 46 |

*completeness was estimated using BUSCO v3.0.2 with lineage data from Cyanobacteria (*8*)

**Table S4: Query sequences used for BLASTP.**

| Filament Protein | Strain | Protein 1D | Functional Reference |
|---|---|---|---|
| CyDiv | *Anabaena* sp. PCC 7120 | WP_010996476.1 | (*9*) |
| HetR | *Anabaena* sp. PCC 7120 | BAB74038 | (*10*) |
| SepJ | *Anabaena* sp. PCC 7120 | WP_010996494.1 | (*11*) |
| Sep1 | *Anabaena* sp. PCC 7120 | WP_044523149.1 | (*3*) |
| HetZ | *Anabaena* sp. PCC 7120 | BAB77623.1 | (*12*) |
| FraE | *Anabaena* sp. PCC 7120 | WP_096637145.1 | (*13, 14*) |
| PatU3 | *Raphidiopsis brookii* D9 | WP_009343449.1 | (*15*) |
| HglK | *Anabaena* sp. PCC 7120 | WP_010994987.1 | (*16*) |
| FraC | *Anabaena* sp. PCC 7120 | WP_010996548.1 | (*13, 14, 17*) |
| FraD | *Anabaena* sp. PCC 7120 | WP_010996549.1 | (*13, 14*) |

**Table S5:** Experimentally-characterised functions of 11 proteins in multicellular Cyanobacteria. Their evolutionary history and distribution among a broad range of strains are modelled in this study because most of the information on their function derives from analyses of model species (often *Anabaena* sp. PCC 7120 from M.S. IV).

| Protein | Function | Citation(s) |
|---|---|---|
| FraC | Aids the localisation and assembly of septal junctions between cells. As a result, it maintains fast intercellular communication and long filaments in nitrogen-free medium. | (*13, 14, 17-19*) |
| FraD | Plugs septal junctions between cells and localises SepJ to the intercellular septa. As a result, it maintains fast intercellular communication and long filaments in nitrogen-free medium. | (*13, 14, 18, 19*) |
| FraE | Maintains fast intercellular communication and long filaments in nitrogen-free medium. Without this permease, heterocysts do not fix as much nitrogen. | (*13, 14*) |
| HetR | Controls the frequency of differentiated (e.g. heterocysts) and vegetative cells in filaments by regulating the transcription of *hetZ* and *hetP.* | (*10, 12, 20-22*) |
| PatU3 | Regulates the frequency of differentiated (e.g. heterocysts) and vegetative cells in filaments by interacting with HetZ | (*15, 20, 21*) |
| HetZ | Regulates the frequency of differentiated (e.g. heterocysts) and vegetative cells in filaments by regulating the expression of *hetR* and *patS* and interacting with PatU3. | (*12, 20, 21, 23*) |
| SepJ | Interacts with peptidoglycan to maintain fast intercellular communication, quick growth rates, long filaments and the development of specialist cells, such as heterocysts. SepJ is also a key component of the septal junctions between cells and interacts with the divisome during cell division to control the width and number of nanopores in septa between cells. SepJ is distinguished from closely-related DMT permeases by the presence of a permease domain, linker region and coiled coils in each protein. | (*11, 14, 19, 24-32*) |
| SepI | Maintains intercellular communication, nanopore formation, septum size and cell morphology by interacting with other septal proteins (e.g. SepJ) as well as proteins associated with the divisome (e.g. ZipN, SepF and FtsI). | (*3*) |
| HglK | This transmembrane protein localises at the intercellular septa, where it aids the formation of nanopores. Without it, few nanopores form and they are smaller. As a result, intercellular communication slows, and filaments shorten. | (*16*) |

| | | |
|---|---|---|
| CyDiv | Aids localisation of the septum during cell division to create a straight filament without kinks | (*9*) |
| Product of *all2344* | Unknown | (*15*) |

**Table S6: Ancestral deviation values of branches used to root bayesian phylogenetic trees of genes associated with filamentous morphology.** The lower the ancestor deviation value, the more likely the root is to be true, and the higher the ambiguity index, the more uncertainty in the choice of the top two rooting positions (*33*).

| Protein | Minimal Ancestor Deviation | Ancestor Deviation of Root | Ambiguity Index | Alignment Length / amino acids |
|---|---|---|---|---|
| CyDiv | 0.215 | 0.250 | 0.999 | 259 |
| HetR | 0.291 | Same | 0.995 | 306 |
| SepJ | 0.207 | Same | 0.959 | 348 |
| SepI | 0.250 | Same | 0.985 | 195 |
| FraE | 0.229 | Same | 0.999 | 264 |
| HetZ and HetZ related protein | 0.200 | Same | 0.995 | 416 |
| PatU3 | 0.181 | 0.183 | 0.989 | 374 |
| HglK | 0.213 | 0.215 | 0.990 | 742 |
